# Altered spawning seasons of Atlantic salmon broodstock transcriptionally and epigenetically influence cell cycle and lipid-mediated regulations in their offspring

**DOI:** 10.1101/2024.02.03.578741

**Authors:** Takaya Saito, Marit Espe, Maren Mommens, Christoph Bock, Jorge M.O. Fernandes, Kaja H. Skjaerven

## Abstract

Manipulating spawning seasons of Atlantic salmon (*Salmo salar*) is a common practice to facilitate year-round harvesting in salmon aquaculture. This process involves adjusting water temperature and light regime to control female broodstock maturation. However, recent studies have indicated that altered spawning seasons can significantly affect the nutritional status and growth performance of the offspring. Therefore, gaining a deeper understanding of the biological regulations influenced by these alterations is crucial to enhance the growth performance of fish over multiple generations.

In this study, we investigated omics data from four different spawning seasons achieved through recirculating aquaculture systems (RAS) and sea-pen-based approaches. In addition to the normal spawning season in November (sea-pen), three altered seasons were designated: off-season (five-month advance, RAS), early season (two-month advance, sea-pen), and late season (two-month delay, sea-pen). We conducted comprehensive gene expression and DNA methylation analysis on liver samples collected from the start-feeding larvae of the next generation.

Our results revealed distinct gene expression and DNA methylation patterns associated with the altered spawning seasons. Specifically, offspring from RAS-based off-season exhibited altered lipid-mediated regulation, while those from sea-pen-based early and late seasons showed changes in cellular processes, particularly in cell cycle regulation when compared to the normal season. The consequences of our findings are significant for growth and health, potentially providing information for developing valuable tools for assessing growth potential and optimizing production strategies in aquaculture.

**Author Summary:** This study examines the impact of manipulating Atlantic salmon spawning seasons in aquaculture on genetic and epigenetic regulations in their offspring. Manipulating water temperature and light cycles during broodstock rearing allows for year-round harvesting. In aquaculture, altering spawning seasons is a common practice; however, recent research suggests that these changes affect the nutritional status and growth of offspring. To understand these effects at the molecular level, we analysed transcriptomic and epigenetic data from salmon offspring born to broodstock with four different spawning seasons. Our results reveal distinct transcriptomic and epigenetic patterns associated with altered seasons, influencing lipid metabolism and cell cycle regulation. These findings hold significant implications for improving aquaculture practices, potentially enhancing growth performance and nutritional quality of farmed salmon across generations as temperature and light are crucial abiotic factors influencing the next generation. Furthermore, our study potentially contributes to understanding the impact of climate change on the spawning behaviour of aquatic vertebrates, particularly in relation to increasing global ocean temperatures and the resulting changes in photoperiod as species migrate northward in the Northern Hemisphere, experiencing longer daylight in summer and shorter daylight in winter.

## Introduction

The Atlantic salmon (*Salmo salar*) follows an anadromous life cycle, beginning in freshwater, migrating to seawater for most of its life, and returning to freshwater to spawn. In salmon aquaculture, the timing of spawning is adjusted to ensure year-round harvesting availability. This manipulation involves controlling abiotic factors, such as water temperature and photoperiod, to influence the spawning timing (Pankhurst and King, 2010; Taranger et al., 2010). Additionally, apart from the traditional open sea-based cages, a land-based approach utilising recirculating aquaculture systems (RAS) is employed to more accurately replicate the salmon’s natural lifecycle, including both saltwater gonad maturation and final freshwater maturation periods (Good and Davidson, 2016). Despite the ongoing and successful practices of spawning season alterations in salmon aquaculture, recent studies have reported that these practices can impact the nutritional status of broodstock, leading to negative effects on the growth performance in the next generation (Skjærven et al., 2022; Skjaerven et al., 2020).

While various nutrients influence the health condition and growth performance of Atlantic salmon (Council, 2011; Hamre et al., 2016; Hemre et al., 2016), altered spawning seasons have appeared to affect the status of multiple metabolites and nutrients including micronutrients, amino acids, and lipid classes (Skjærven et al., 2022; Skjaerven et al., 2020). These nutrients are also closely associated with vital biological functions necessary for growth, such as one carbon (1C) metabolism, the citric acid cycle, and the Cahill cycle. Multiple key micronutrients, including vitamin B12, folate, vitamin B6, and methionine, are integrated with the 1C metabolism (Danchin et al., 2020; Lyon et al., 2020), which regulates cellular methylation potential through S-adenosylmethionine (SAM) and S-adenosylhomocysteine (SAH), critical for various cellular processes (Ducker and Rabinowitz, 2017). The citric acid cycle enables the catabolism of carbohydrates, fats, and amino acids, releasing stored energy crucial for growth and development (Akram, 2014), while the Cahill cycle facilitates the transport of amino groups and carbons between muscle and liver (Felig, 1973), both playing essential roles in various developmental processes. Amino acids are essential for growth and tissue development, in addition to being the building blocks of proteins (Wu, 2010), while lipids serve as major energy reserves and contribute to the formation of cell membranes, also supporting various developmental processes (Fahy et al., 2011).

Beyond the importance of nutrients, understanding genetic influences provides invaluable insights from a molecular biology perspective. Recent studies have revealed that alterations in micronutrient levels within the feed can have transcriptomic effects on various biological pathways in salmon, including lipid metabolism, protein synthesis, and post-transcriptional regulation (Saito et al., 2023; Saito et al., 2021). Moreover, an additional layer of information comes from the analysis of DNA methylation, which supports to reveal important epigenetic regulations. DNA methylation, as an epigenetic regulatory process, influences various cellular mechanisms, including cell stability, differentiation, and development, potentially in response to environmental stimuli. (Amenyah et al., 2020; Moore et al., 2013). Since DNA methylation is inherited during cell replication, it can result in long-term changes in epigenetic regulation (Goll and Bestor, 2005; Moore et al., 2013).

The present study investigates the transcriptomic and epigenetic effects on offspring resulting from four different spawning seasons. These seasons include the natural sea-pen-based spawning in November, an accelerated RAS-based spawning occurring five months earlier, and two variations of sea-pen-based spawning: one with a two-month advance and another with a two-month delay relative to the natural spawning season. The main analysis of the study focuses on omics analysis of liver samples collected from start-feeding larvae at the end of the endogenous feeding phase. The liver serves as a key target organ for various metabolic reactions crucial to growth, and using start-feeding larvae helps eliminate the influences of exogenous feeding. To uncover both transcriptomic and epigenetic impacts comprehensively, we employed RNA-sequencing (RNA-seq) for gene expression and reduced-representation bisulfite sequencing (RRBS) for DNA methylation analysis. Our findings suggest that gene expression and DNA methylation data can serve as powerful tools for assessing current and future growth performance. These tools could potentially benefit a wide range of nutritional programming studies and enhance our understanding of how aquatic species adapt to climate change.

## Methods and Materials

### Ethical considerations

The experiment adhered to the ARRIVE guidelines for design and reporting. Broodstock and offspring used in the study were obtained from AquaGen’s commercial production facility of Atlantic salmon at their breeding station in Kyrksæterøra, Norway.

As the samples were collected at the breeding station, the conditions and protocols applied in this experiment were identical to those used in commercial production, including certain confidential information protected under commercial interests. Throughout the sampling procedures for broodstock and larvae, anaesthetics were administered by the supplier’s instructions and compliance with both Norwegian and European legislation on animal research.

Formal approval from the Norwegian Animal Research Authority (NARA) was not required for this experiment. This exemption is because the experimental conditions fall under practices recognized as commercial animal husbandry, thereby exempting them from the European Convention on the protection of animals used for scientific purposes (2010/63/EU), specifically under Article 5d. Additionally, the Norwegian ethics board also approved the experiment by the Norwegian regulation on animal experimentation, § 2, 5a, d, for activities classified as “non-experimental husbandry (agriculture or aquaculture)” and “procedures in normal/common breeding and husbandry”.

### Experimental design and sampling

As the details of experimental design and sampling methods have been described elsewhere (Skjærven et al., 2022; Skjaerven et al., 2020), we only present summarized descriptions here. The study utilized broodstock and offspring obtained from AquaGen’s breeding station in Kyrksæterøra, Norway. The female broodstock, sourced from the 15th generation, covered four spawning seasons: off-season, early, normal, and late seasons.

The off-season broodstock were reared in a land-based recirculating aquaculture system (RAS) in brackish (12‰ salinity), while the broodstock from the early, normal, and late seasons were raised in three open sea-based net pens with ambient photoperiod and temperatures. All broodstock were fed to satiation with broodstock feed (EWOS Breed 3500) until transferred to freshwater. In freshwater, the normal season broodstock matured naturally under ambient photoperiod and temperatures, whereas artificial abiotic factors like temperature and light were applied to regulate maturation for the other spawning seasons. Specifically, while the normal season broodstock were maintained at 6 °C with an 8-hour daylight cycle upon transfer to freshwater tanks in August, the early season broodstock were maintained by following the protocol outlined by Naeve and colleagues (Naeve et al., 2018). Details about the temperature and photoperiod used to postpone maturation in the off-season and late season groups are proprietary information owned by AquaGen AS. The off-season and early season broodstock were subjected to five– and two-month advance maturation, respectively, while the late season broodstock were subjected to a two-month delay. The fish received no further feeding after the transfer to freshwater. The maturation process in freshwater lasted for 109, 163, 120, and 166 days for off-season, early, normal, and late season broodstock, respectively, before fertilization to obtain the next generation (Supplementary Table S1).

From each spawning group, five females from separate tanks were sacrificed using Benzoak (200 mg/L, ACD Pharmaceuticals AS). After stripping, each female’s growth measures, including body weights (n=5 per spawning season, randomly selected), were recorded. Liver samples were dissected and flash frozen in liquid N2 for nutrient analysis. All oocytes were fertilised with cryopreserved sperm from two pooled males. Newly fertilized eggs (2 d°) were flash-frozen in liquid N2 for nutrient analysis (Supplementary Table S1).

At the end of the endogenous feeding phase (start-feeding larvae 979–994 d°; Supplementary Table S1), 36 larvae from each broodstock were randomly selected, euthanised with an overdose of buffered tricane methanesulfonate (MS222, Pharmaq, Norway), and body weights (n=180 per spawning season) were recorded. Additionally, 40 liver samples (two from each broodstock fish) were dissected from larvae and flash frozen in liquid N2 and used for either RNA or DNA analysis.

### Growth measure and nutrient analysis

We collected the data on body weights for broodstock and larvae from previous studies (Skjærven et al., 2022; Skjaerven et al., 2020). In addition, we obtained the data on selected nutrients and metabolites from the same studies, specifically vitamin B6 & B12, folate, S-adenosylmethionine (SAM), S-adenosylhomocysteine (SAH), cholesterol, the sum of six different lipids, glutamate, L-serine, L-lysine, L-alanine, L-glutamine, urea, and B-alanine (Skjærven et al., 2022; Skjaerven et al., 2020). To analyse the data from all four spawning seasons, we performed ANOVA followed by Tukey’s HSD test.

### DNA and RNA extraction

RNA extraction from 20 larvae livers (n=5 for each spawning season) was carried out using the BioRobot EZ1 and EZ1 RNA Universal Tissue kit (Qiagen), followed by DNase treatment using the Ambion DNA-free DNA removal kit (Invitrogen, USA) according to their respective protocols. The RNA quantity was assessed using the NanoDrop ND-1000 Spectrophotometer (Nanodrop Technologies) and the Agilent 2100 Bioanalyzer with the RNA 6000 Nano LabChip kit (Agilent Technologies). For DNA isolation from 20 larvae livers (n=5 for each spawning season), the DNeasy Blood & Tissue Kit (Qiagen, Cat. No. #69506) was used as per the manufacturer’s protocol. Liver samples underwent RNase A treatment (provided by the Qiagen kit, 50ng/µL, 10 min at room temperature) followed by proteinase K treatment (New England Biolabs, #8102S 20µg/µL, 1.5 h at 55°C). DNA was eluted in Milli Q water. DNA quantification was performed using the Qubit High Sensitivity Assay (Life Technologies #Q32854). Detailed methods for both DNA and RNA extraction were described elsewhere (Saito et al., 2021).

### Library preparation for RNA-seq and RRBS

Liver samples were sequenced at two different sequencing facilities for specific analyses. For RNA-seq, the liver samples were processed at the DeepSeq sequencing facility at Nord University, Bodø, Norway, and the library preparation was conducted using the NEBNext Ultra II Directional RNA Library Prep Kit for Illumina (New England Biolabs). Sequencing was performed on the NextSeq500 machine (Illumina). For RRBS, the liver samples were processed at the CeMM Biomedical Sequencing Facility, Vienna, Austria. The genomic DNA was extracted, digested by MspI, and bisulfite-converted before library preparation. The RRBS libraries were sequenced on Illumina HiSeq 3000/4000 instruments. Detailed methods of the library preparation for RNA-seq and RRBS were described elsewhere (Saito et al., 2021).

### Atlantic salmon genome and genomic annotation

The reference genome (ICSASG version 2) and RefSeq data (version 100) for gene annotation were downloaded from the NCBI web site (https://www.ncbi.nlm.nih.gov/assembly/GCF_000233375.1). In cases where gene symbols were either outdated or unavailable from RefSeq data (version 100), gene symbols from a newer version of RefSeq data (version 102) and UniProt (Bateman et al., 2022) were used. The genome regions were annotated into intron, exon, three promoter regions, and flanking regions. Promoter regions were divided into three categories based on their distance from transcription start sites: P250 (1-250 bp), P1K (251-1000 bp), and P5K (1001-5000 bp). Flanking regions were defined as 5000 upstream from P5K and 10000 downstream from the 5’ end of the gene.

### Pre-processing of high throughput sequencing

For both RNA-seq and RRBS data, the same pre-processing procedures were followed as described in a previous study (Saito et al., 2021). This involved trimming the reads using Cutadapt (Martin, 2011) and Trim Galore! (Barbraham Institute), aligning the trimmed reads to the reference genome using STAR (Dobin et al., 2013) for RNA-seq and Bismark/Bowtie 1 (Krueger and Andrews, 2011; Langmead et al., 2009) for RRBS with their default parameters. The mapped RNA-seq reads were quantified using featureCounts (Liao et al., 2014), while the mapped RRBS reads were processed by Bismark (Krueger and Andrews, 2011) for methylation calling and CpG site extraction. The samples were divided into four spawning season groups for further analysis: off-season, early, normal, and late seasons, and clustering analysis was performed using the factoextra package (https://CRAN.R-project.org/package=factoextra). RNA-seq counts were subjected to a variance stabilizing transformation (VST) using DESeq2 (Love et al., 2014) before principal component analysis (PCA).

Raw reading data were uploaded to the repository of the Sequence Read Archive (SRA) on the NCBI web site and available under the accession numbers PRJNA680425 and PRJNA642998 for RNA-seq and RRBS, respectively.

### Differential gene expression analysis

Differentially expressed genes (DEGs) were identified using DESeq2 (Love et al., 2014) with an adjusted p-value cut-off of less than 0.05. P-values were adjusted by the Benjamini-Hochberg procedure (Benjamini and Hochberg, 1995). From the candidate DEGs, those with absolute log fold changes (LFCs) greater than or equal to 1.2 were selected as the final set of DEGs by using the lfcThreshold argument of the results function of DESeq2 (Love et al., 2014). Shrunken LFCs values, calculated by the normal shrinkage method provided by DESeq2 (Love et al., 2014), were used for both heatmaps showing functional annotation results and scatter plots showing merged results between DNA methylation and gene expression differences.

### Functional annotation with KEGG

Over-representation analysis (ORA) was performed on the Kyoto Encyclopedia of Genes and Genomes (KEGG) database (Kanehisa and Goto, 2000a) using clusterProfiler (Yu et al., 2012). Gene lists of DEGs identified from the three pair-wise comparisons against the normal season were used as input, and enriched pathways were determined based on adjusted p-values less than 0.05. P-values were adjusted by the Benjamini-Hochberg procedure (Benjamini and Hochberg, 1995).

### Methylation rate analysis

Mapped RRBS reads were filtered by methylKit (Akalin et al., 2012), discarding reads with coverage less than or equal to 10 and above the 99.9th percentile. Additional RRBS data from two other studies were obtained for post-smolt and harvesting stages (Saito et al., 2023; Saito et al., 2021) and underwent the same pre-processing procedures to compare the distributions of methylation rates. Raw data of these RRBS sequencing data can be found in SRA on the NCBI site under the accession numbers of PRJNA680423 and PRJNA628740 for post-smolt and harvesting stages, respectively.

### Differential methylation analysis

Differentially methylated CpG sites (DMCs) were identified by methylKit (Akalin et al., 2012), and Q-values were calculated using the sliding linear model (SLIM) method (Wang et al., 2011). CpGs were identified as DMCs when Q-values were less than 0.01 and methylation differences were greater than or equal to 15%.

To filter out DMCs that were strongly affected by altered spawning seasons, we used the counts of DMCs per gene in two promoter regions: P250 and P1K. Genes were sorted by the count of DMCs in a descendant order, separately for P250 and P1K, and genes that contained at least three DMCs were selected for further analysis.

### Bioinformatics analysis

In-house R and Python scripts with Snakemake (Koster and Rahmann, 2012) were utilized for high-throughput sequence analysis, basic statistical analysis, and figure generation.

## Results

### Altered spawning seasons impacted the weight and nutritional status of offspring

The present study examines the effects on gene expression and DNA methylation patterns in the liver of offspring hatched from four different spawning seasons: an off-season based on a recirculating aquaculture system (RAS) and three sea-pen based seasons (early, normal, and late) (Fig. 1a). Off-season spawning occurred in June, early season in September, and late season in January, relative to the normal spawning months of November. (see detailed dates and degree days in Supplementary Table S1). The growth performance and nutritional status of the broodstock and offspring have already been assessed and discussed elsewhere (Skjærven et al., 2022; Skjaerven et al., 2020). This section provides a summary of previous research, emphasizing the weight and the status of selected nutrients, while the primary focus of the present study is on the omics analysis.

**Fig. 1.**
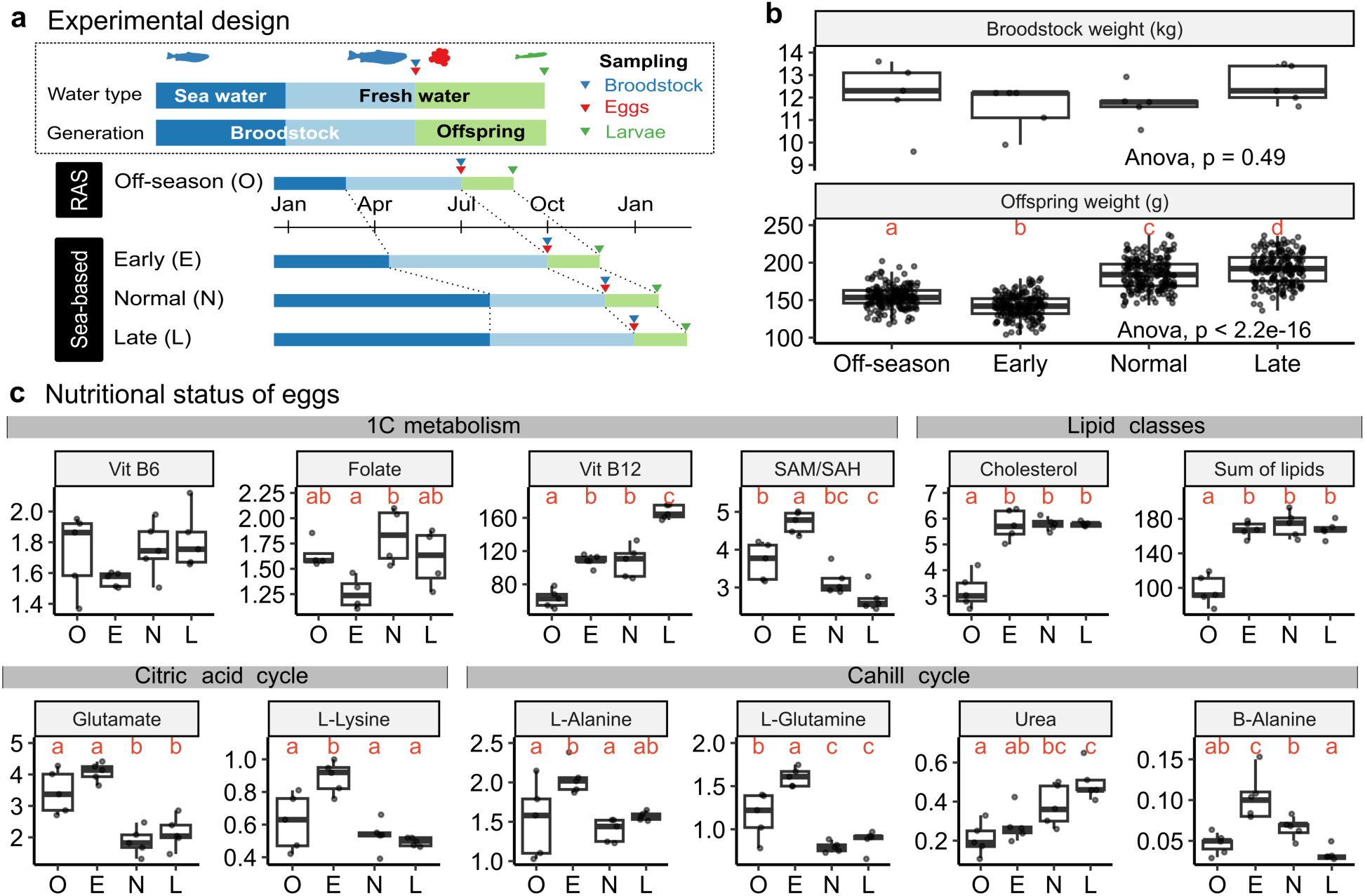
Experimental design, body weights, and nutritional status across spawning seasons. **a**) Schematic diagram illustrating four distinct spawning seasons: off-season (O), early (E), normal (N), and late (L). The off-season was conducted in RAS, while others were in a sea-based environment. Broodstock were transferred from seawater (blue bars) to freshwater (light blue bars) before spawning. Offspring were raised in freshwater until sampling (green bars). Triangles indicate sampling points: broodstock (blue), eggs (red), and larvae (green). **b)** Boxplots showing body weights of broodstock (n=5) and larvae (n=180) per season. Red letters represent Tukey’s HSD compact letter display after ANOVA tests (p<0.05). **c)** Boxplots of 12 nutrients and metabolites in egg samples. Nutrients and metabolites are categorised into four groups: 1C metabolism, lipid classes, citric acid cycle, and Cahill cycle. Units: 1C metabolism group (Vitamin B6: mg/kg ww, Folate: mg/kg ww, Vitamin B12: µg/kg ww); remaining groups: lipid classes (mg/g ww), citric acid cycle (µmol/g ww), Cahill cycle (µmol/g ww). Red letters represent Tukey’s HSD compact letter display after ANOVA tests (p<0.05).

Broodstock weights showed no significant differences at spawning (upper part of Fig. 1b), but offspring weights at the larvae stage (979 and 994 degree days) showed noticeable variation (lower part of Fig. 1b). Specifically, off-season and early season larvae had lower weights compared to normal and late seasons (Supplementary Table S2). This trend was consistent with the observations in egg sizes, where eggs from off-season and early-season exhibited smaller sizes compared to the other two seasons (see the number of eggs per liter in Supplementary Table S1).

Altered spawning seasons widely impacted the nutritional status of offspring eggs, particularly regarding 1C metabolism, lipid classes, citric acid cycle, and Cahill cycle (Skjærven et al., 2022; Skjaerven et al., 2020), but with varying patterns across nutrients and metabolites (summarised in Fig. 1c and Supplementary Tables S3-S6). Previous studies also reported that altered spawning seasons affected the nutritional status of the broodstock in addition to their offspring (Skjærven et al., 2022; Skjaerven et al., 2020). Thus, altered spawning seasons significantly influenced the nutritional status of broodstock without impacting their weights, however, subsequently affecting offspring nutritional status and growth potential of their offspring.

### Early and late spawning seasons exhibited similar gene expression patterns when compared to the normal season

To investigate the impact of different spawning seasons, we performed gene expression analysis on 20 liver samples, which were divided into four groups (n=5). We compared gene expression from the normal season to that from the other three altered spawning seasons (detailed statistics for each sample are provided in Supplementary Table S7).

Principal component analysis (PCA) differentiated the off-season from the early and normal seasons when considering all mapped genes (left part of Fig. 2a). However, when focusing on the top 500 genes with high variances, the off-season moderately deviated from the early season but not notably from the normal season (right part of Fig. 2a). In both cases, the late season overlapped partially with all other seasons.

**Fig. 2.**
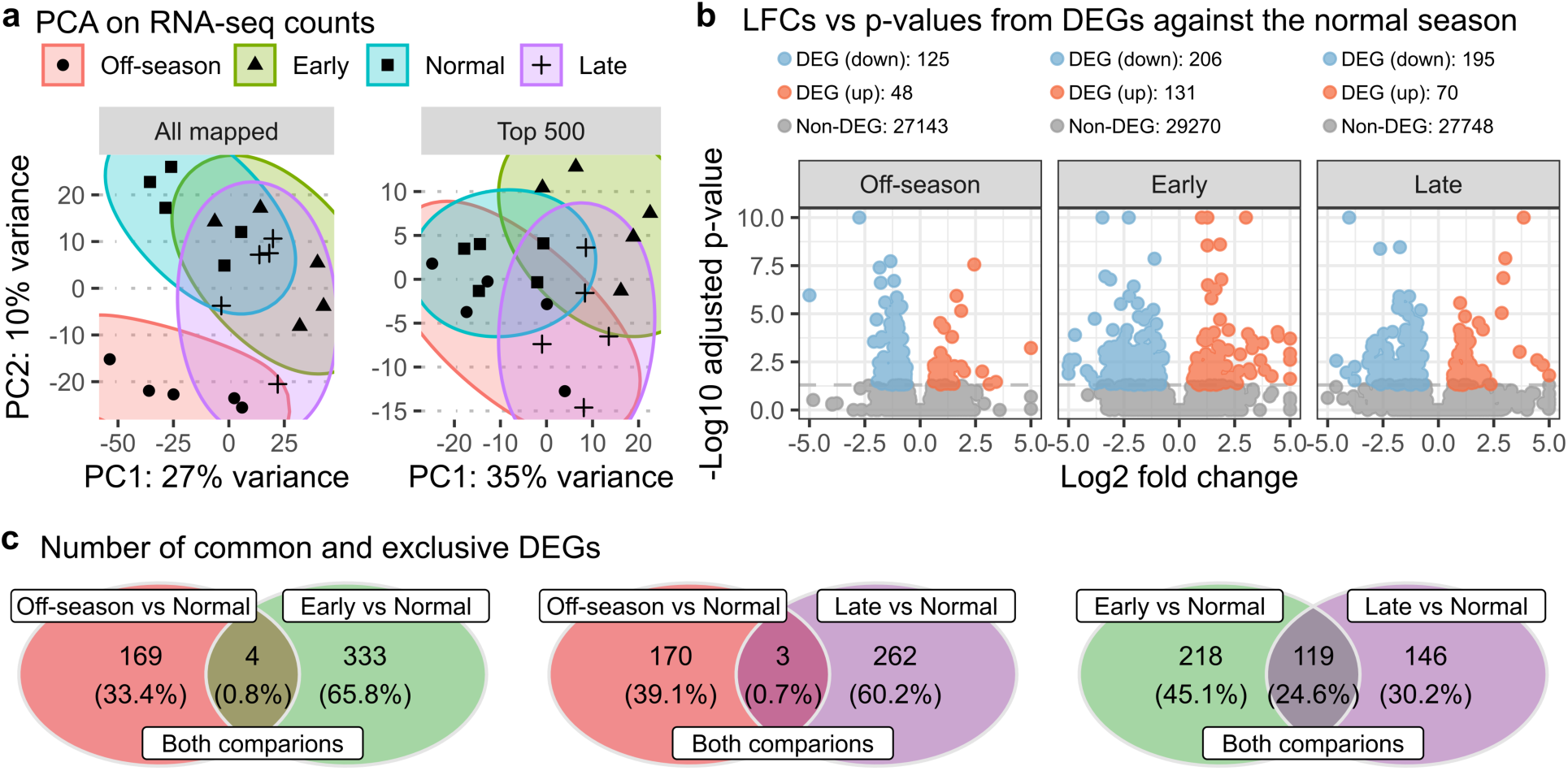
Gene expression analysis comparing altered spawning seasons to the normal season. **a**) PCA plots displaying clustering patterns of four spawning seasons – off-season (red), normal (green), early (blue), and late (purple) – using 20 RNA-seq samples. “All mapped” plot includes 45153 genes, while “Top 500” includes top 500 genes with high variance. **b)** Volcano plots showing differential expression analysis results for three pairwise comparisons: off-season vs. normal (Off-season), early vs. normal (Early), and late vs. normal (Late). Dots represent down-regulated DEGs (blue), up-regulated DEGs (red), and non-DEGs (grey). The counts of DEG and non-DEGs were provided in the legend. **c)** Venn diagrams illustrating overlaps of DEGs identified between two comparisons, indicating common DEGs as well as those exclusive to a single comparison.

In three pairwise comparisons for differential expression analysis against the normal season (off-season vs. normal, early vs. normal, and late vs. normal), we identified 173, 337, and 265 differentially expressed genes (DEGs), respectively (Fig. 2b). All comparisons revealed a higher number of down-regulated genes than up-regulated genes. Additionally, Venn diagrams indicated that the DEGs from the off-season vs. normal comparison had very few overlapping genes (<1%) with those from the other two comparisons (depicted in the first and second diagrams in Fig. 2c). On the other hand, early vs. normal and late vs. normal comparisons shared approximately 25% of the DEGs (depicted in the third diagram in Fig. 2c). These results suggest that the expression pattern of the off-season differed from both the early and late seasons when compared to the normal season.

### Altered spawning seasons affected various biological pathways related to metabolism, cellular processes, and organismal systems

To explore the impact of altered spawning seasons on biological pathways, we conducted functional analysis on DEGs using the Kyoto Encyclopedia of Genes and Genomes (KEGG) database. Over-representation analysis (ORA) identified a total of 11 enriched KEGG pathways, which were classified into three functional categories: metabolism, cellular processes, and organismal systems, when comparing the three altered spawning seasons against the normal season (Fig. 3).

**Fig. 3.**
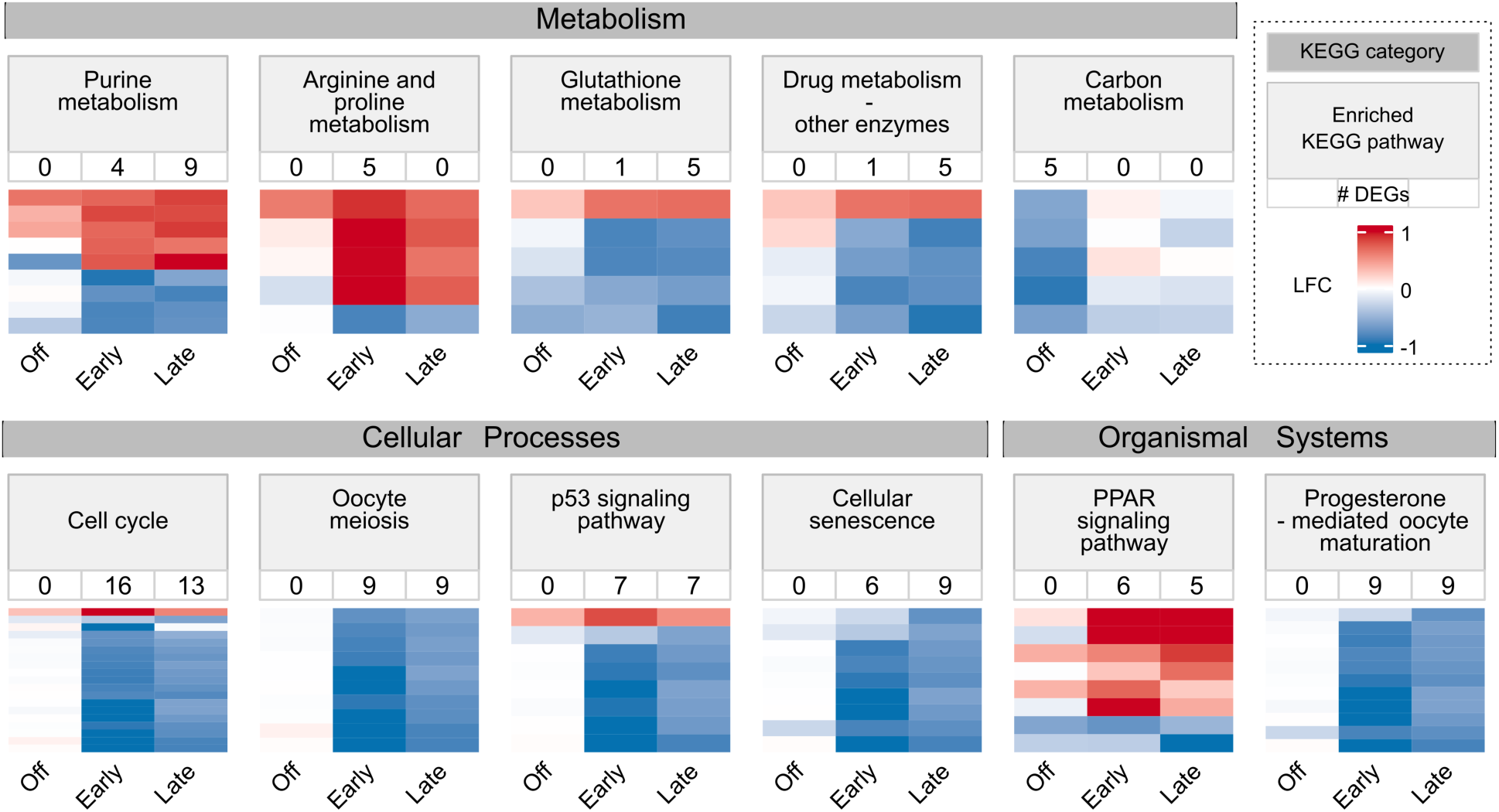
Enriched KEGG pathways for DEGs identified in three pair-wise comparisons. Heatmaps showing log fold changes (LFCs) of genes associated with 11 KEGG pathways enriched through over-representation analysis. LFCs underwent normal shrinkage transformation. Genes were included if identified as DEGs in at least one of the three comparisons: off-season vs normal (Off), early vs normal (Early), and late vs normal (Late). The number below each pathway represents DEG count. KEGG pathways are categorized into metabolism, cellular processes, and organismal systems groups.

The off-season vs. normal comparison identified only one enriched pathway, carbon metabolism (Supplementary Table S8). All five DEGs associated with carbon metabolism showed down-regulation (Fig. 3).

The early vs. normal comparison revealed six enriched pathways: one in metabolism, three in cellular processes, and two in organismal systems (Supplementary Table S9). Metabolism-related pathways exhibited both up– and down-regulation, while most genes in cellular processes showed significant down-regulation. In organismal systems, PPAR signalling showed up-regulation, while progesterone-mediated oocyte maturation showed down-regulation (Fig. 3).

The late vs. normal comparison identified nine enriched pathways: three in metabolism, four in cellular processes, and two in organismal systems (Supplementary Table S10). Remarkably, this comparison shared five enriched pathways with the early vs. normal comparison, indicating similar impacts on biological pathways between them. Moreover, these pathways displayed consistent expression patterns in terms of up– and down-regulation between early and late seasons. Notably, genes associated with cellular processes exhibited strong down-regulation for both seasons (Fig. 3).

In summary, the off-season appeared to down-regulate carbon metabolism, while the early and late seasons primarily had strong suppressive effects on pathways associated with cellular processes.

### Altered spawning seasons had an impact on the expression of genes in a similar manner

To further investigate the genes commonly affected by multiple spawning seasons, we used overlapping DEGs (represented as intersections of Venn diagrams in Fig. 2c) by applying two filtering steps: selecting DEGs found in at least two comparisons and choosing the top five DEGs from each comparison based on adjusted p-values (Supplementary Table S11). This filtering approach revealed a total of 13 genes (Table 1). Among them, two genes (gene IDs: 106612264 and 106601246) lacked corresponding gene names and symbols. All genes, except *pc* and *pitpnm2*, consistently displayed up-or down-regulation across all spawning seasons, suggesting that all altered spawning seasons had a similar effect on this specific set of genes when compared against the normal season (Table 1).

**Table 1.**
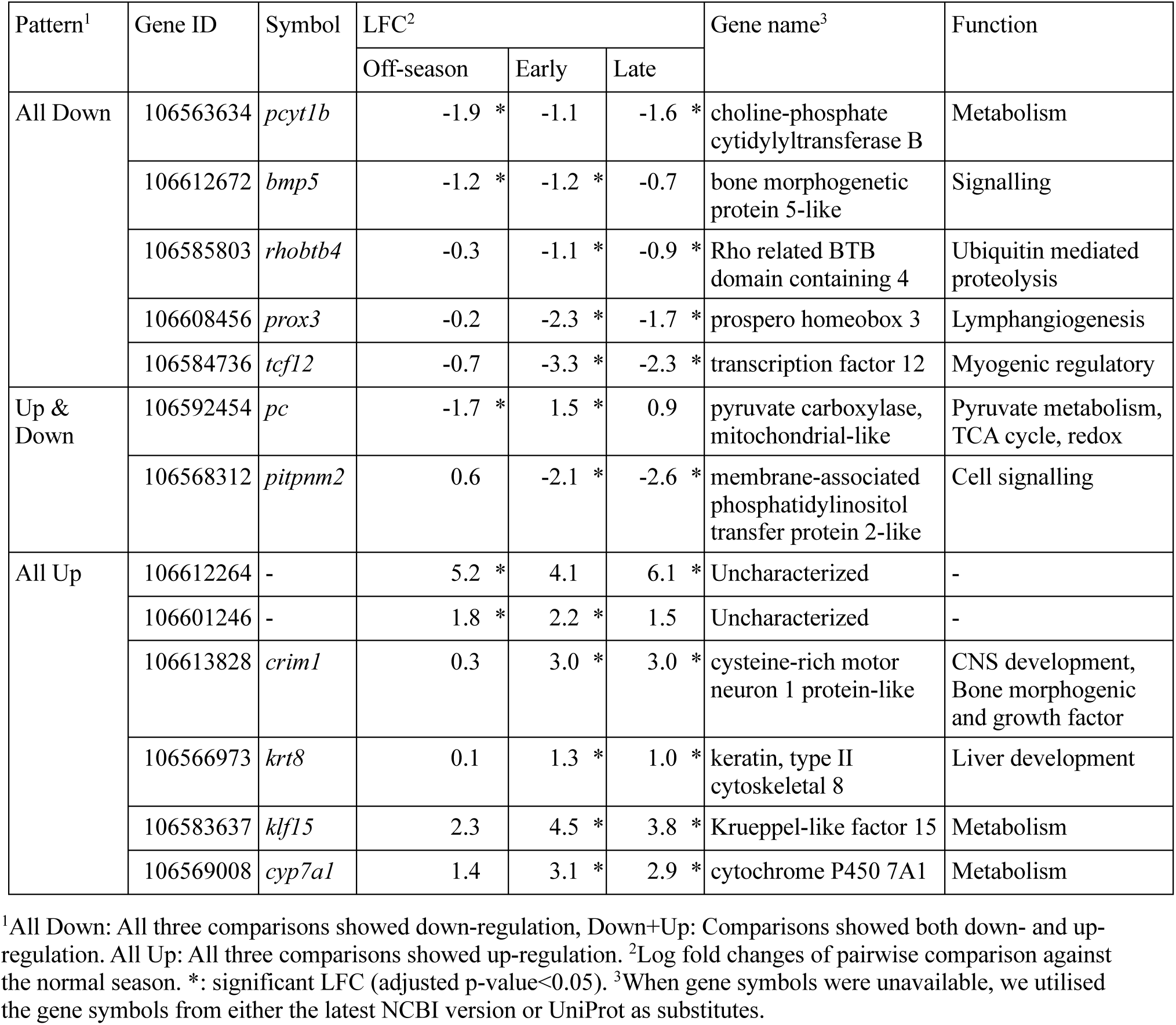
List of top five DEGs identified in at least two pairwise comparisons.

| Pattern<sup>1</sup> | Gene ID | Symbol | LFC<sup>2</sup> |  |  | Gene name<sup>3</sup> | Function |
| --- | --- | --- | --- | --- | --- | --- | --- |
|  |  |  | Off-season | Early | Late |  |  |
| All Down | 106563634 | *pcyt1b* | -1.9 * | -1.1 | -1.6 * | choline-phosphate cytidyltransferase B | Metabolism |
|  | 106612672 | *bmp5* | -1.2 * | -1.2 * | -0.7 | bone morphogenetic protein 5-like | Signalling |
|  | 106585803 | *rhobtb4* | -0.3 | -1.1 * | -0.9 * | Rho related BTB domain containing 4 | Ubiquitin mediated proteolysis |
|  | 106608456 | *prox3* | -0.2 | -2.3 * | -1.7 * | prospero homeobox 3 | Lymphangiogenesis |
|  | 106584736 | *tcf12* | -0.7 | -3.3 * | -2.3 * | transcription factor 12 | Myogenic regulatory |
| Up & Down | 106592454 | *pc* | -1.7 * | 1.5 * | 0.9 | pyruvate carboxylase, mitochondrial-like | Pyruvate metabolism, TCA cycle, redox |
|  | 106568312 | *pitpnm2* | 0.6 | -2.1 * | -2.6 * | membrane-associated phosphatidylinositol transfer protein 2-like | Cell signalling |
| All Up | 106612264 | - | 5.2 * | 4.1 | 6.1 * | Uncharacterized | - |
|  | 106601246 | - | 1.8 * | 2.2 * | 1.5 | Uncharacterized | - |
|  | 106613828 | *crim1* | 0.3 | 3.0 * | 3.0 * | cysteine-rich motor neuron 1 protein-like | CNS development, Bone morphogenic and growth factor |
|  | 106566973 | *krt8* | 0.1 | 1.3 * | 1.0 * | keratin, type II cytoskeletal 8 | Liver development |
|  | 106583637 | *klf15* | 2.3 | 4.5 * | 3.8 * | Krueppel-like factor 15 | Metabolism |
|  | 106569008 | *cyp7a1* | 1.4 | 3.1 * | 2.9 * | cytochrome P450 7A1 | Metabolism |

<sup>1</sup>All Down: All three comparisons showed down-regulation, Down+Up: Comparisons showed both down- and up-regulation. All Up: All three comparisons showed up-regulation. <sup>2</sup>Log fold changes of pairwise comparison against the normal season. \*: significant LFC (adjusted p-value<0.05). <sup>3</sup>When gene symbols were unavailable, we utilised the gene symbols from either the latest NCBI version or UniProt as substitutes.

These genes can be classified into three categories by relevant biological functions: metabolism (*pcyt1b*, *pc*, *klf15*, and *cyp7a1*) (Cappel et al., 2019; Kanehisa and Goto, 2000b; Takashima et al., 2010), tissue development (*prox3*, *crim1*, and *krt8*) (Del Giacco et al., 2010; Kolle et al., 2000; Ku et al., 2007; Scott et al., 2007), and information processing, which can be further classified into three sub categories as signalling (*bmp5* and *pitpnm2*) (Garner et al., 2012; Kanehisa and Goto, 2000b), transcription (*tcf12*) (Hu et al., 1992), and post-translation (*rhobtb4*) (Kanehisa and Goto, 2000b). As these categories are typically associated with growth, all three altered spawning seasons affected a certain set of genes associated with growth regulation in a similar manner, despite PCA plots (Fig. 2a) and Venn diagrams (Fig. 2c) showing a distinct gene expression pattern of the off-season.

### Overall methylation rates increased until the harvesting stage within and around gene bodies, except near transcription start sites

We conducted DNA methylation analysis on 20 liver samples from larvae across four different spawning seasons (n=5; detailed statistics in Supplementary Table S12) to investigate the epigenetic effects of altered spawning seasons on DNA methylation.

Before analysing the effects of altered spawning seasons, we used RRBS data from two independent studies on liver samples to investigate DNA methylation changes across developmental stages. Specifically, we compared our RRBS data from the larvae stage with the post-smolt stage (approximately 31 weeks) and the harvest stage (around 45 weeks) (Saito et al., 2023; Saito et al., 2021). The post-smolt samples were divided into three groups with different 1C metabolism nutrient levels (Ctrl, 1C+, and 1C++) according to their experimental diets, while the harvest samples were divided into three groups with varying micronutrient levels (L1, L2, and L3) based on their experimental diets. We compared these six groups from post-smolt and harvest developmental stages with our larvae stage samples, which were collected across four spawning seasons (Fig. 4a).

**Fig. 4.**
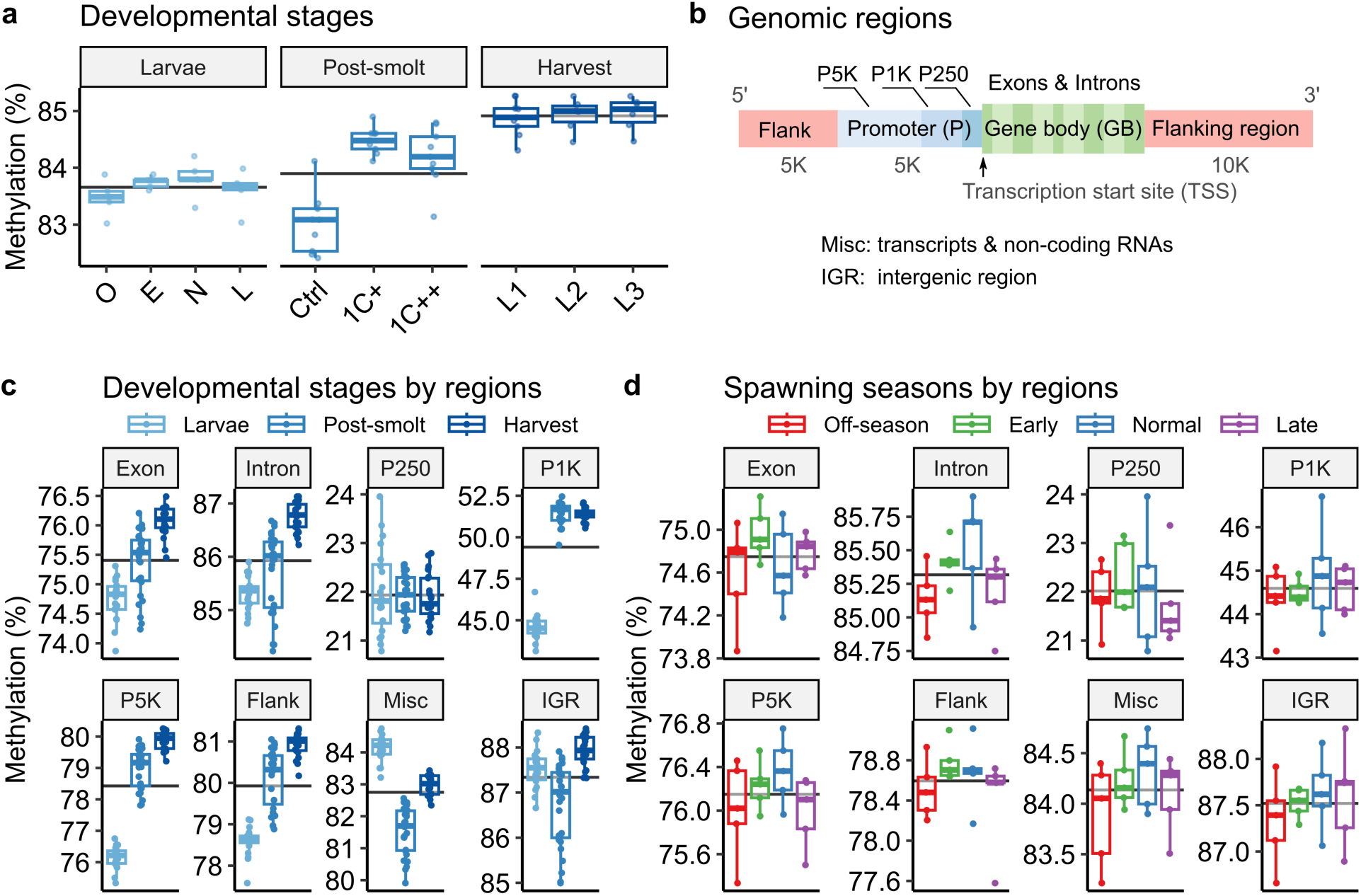
Distribution of methylated CpG sites across genomic regions. Box plots displaying liver DNA methylation status during the lifespan of Atlantic salmon; data from the present study (larvae, light blue) and two other studies: post-smolt (blue) and harvest (dark blue). In the larvae stage, O, E, N, L indicate off-season, early, normal, and late seasons, respectively. While the post-smolt stage contains three groups with different 1C-metabolism nutrient levels (Ctrl, 1C+, and 1C++), the harvest stage contains three groups with different micronutrient levels (L1, L2, and L3) in their diets. Horizontal lines indicate the average rates for each group. **a)** Box plots showing average methylation rates of the samples from three developmental stages. **b)** Genomic region diagram illustrating three regions: flanking regions (Flank, red), promoters (P, blue), and gene bodies (GB, green). The promoter region is divided into three subregions: P250, P1K, and P5K. The gene body includes exons and introns. **c)** Box plots showing average methylation rates of the samples from three developmental stages across eight genomic regions. **d)** Box plots showing average methylation rates of the samples from four spawning seasons across eight genomic regions.

Notably, the average methylation rates exhibited a consistent increase from larvae to post-smolt to harvest stages: 83.7%, 83.9%, and 84.9% for larvae, post-smolt, and harvest stages, respectively (Kruskal-Wallis, p-value < 0.001, black horizontal lines in Fig. 4a). Within the larvae group, the average rate of the normal group was higher than that of the off-season group, but the difference was not significant. Within the post-smolt group, the Ctrl group showed a significantly lower methylation rate than the other two groups (pair-wise Wilcoxon test with the Holm correction, p-values < 0.05; Supplementary Table S13). The Ctrl group was given lower methionine along with B-vitamins as the 1C metabolism nutrients compared to other groups (1C+ and 1C++) (Saito et al., 2023).

To examine regional DNA methylation patterns, CpG sites were categorised into three genomic regions based on their positions relative to nearby genes: gene body (GB; including exons and introns), promoter (P; including P250, P1K, and P5K), flanking regions, miscellaneous regions (Mics, including various transcripts and non-coding RNA), and intergenic regions (IGR) (Fig. 4b; see Methods and Materials for definitions).

The trend of the rate increment became more noticeable when the rates were examined separately by different genomic regions (Fig. 4c). In most regions, including gene bodies, promoter regions except for P250, and flanking regions, the samples from the harvest stage exhibited significantly higher average rates, followed by the post-smolt samples, and then the larvae samples (pair-wise Wilcoxon test with the Holm correction, p-values < 0.05; Supplementary Table S14). Specifically, within the two promoter regions (P1K and P5K), both the post-smolt and harvest stages showed a significant increase in methylation rates compared to the larvae stage. Moreover, methylation rates were generally lower in P250 (∼22%) and P1K (∼49%) compared to other regions (75% ∼ 88%).

This comparative investigation suggests a potential increase in methylation rates as salmon progress in growth, at least until the harvesting stage, while also noting that methylation rates near transcription start sites (TSSs) tend to remain lower and more stable than in other genomic regions.

### The off-season showed non-significant but consistently lower methylation rates compared to the normal and early seasons

To investigate the regional shift of the methylation rates among four spawning seasons, we applied the same methods used for the comparison analysis between developmental stages. Unlike the result of the developmental stages, spawning seasons exhibited no significant differences in methylation rates when separated by genomic regions (Fig. 4d; pair-wise Wilcoxon test with the Holm correction in Supplementary Table S15). Despite the non-significant differences, the average rates of the off-season samples were lower than those of the normal and early seasons in all regions (Supplementary Table S16). Nonetheless, the average rates of the off-season were higher than those of the late season in the two promoter regions (P250 and P5K) and flanking regions.

### Differential methylation patterns indicated higher dynamic regulation in promoter regions compared to other regions

We conducted six pairwise comparisons to identify CpG sites with differential methylation (Fig. 5a). Unlike the analysis of the RNA-seq data, we considered all six possible pair-wise comparisons to capture a comprehensive pattern of methylation differences with our RRBS data. Remarkably, all comparisons consistently demonstrated similar characteristics, for both the number of differentially methylated CpGs (DMCs) and the balanced ratio of hypo– and hyper-methylation (Fig. 5a).

**Fig. 5.**
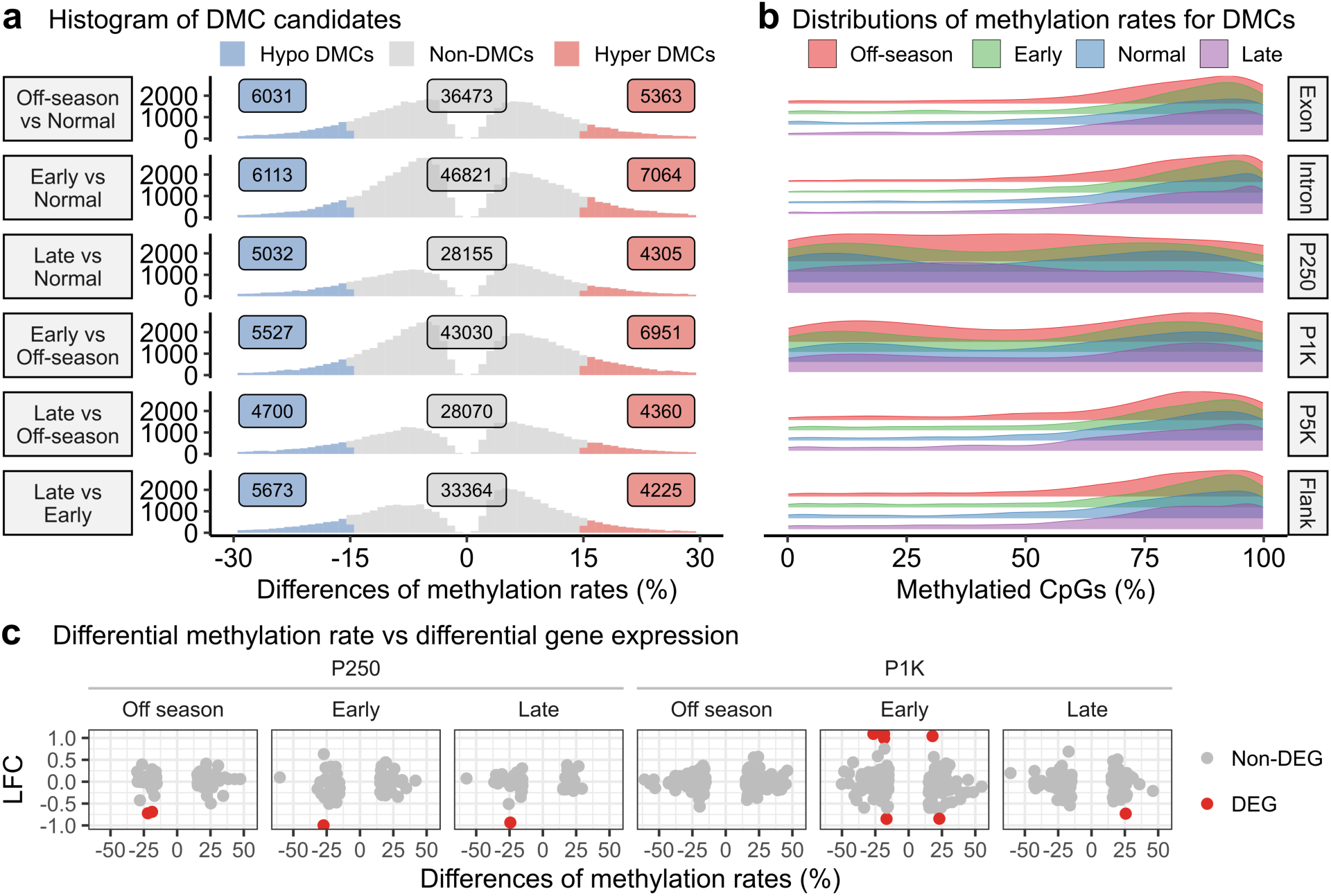
DNA methylation differences and correlations with gene expression. **a**) Histograms displaying distributions of methylation rate differences in six pairwise comparisons. Hypo-methylated DMCs in blue, hyper-methylated DMCs in red, non-DMCs in grey. All CpG sites in plots showed significant methylation differences. **b)** Ridge density plots illustrating distribution patterns of methylated CpG sites across four spawning seasons: off-season (red), early (green), normal (blue), and late (purple). CpG sites were selected if at least one site identified as DMC in any of six comparisons. **c)** Scatter plots showing the relationship between methylation rate differences and log fold changes (LFCs) of gene expression. DMCs were selected from three comparisons: off-season vs. normal (Off-season), early vs. normal (Early), and late vs. normal (Late), and also selected based on two regions, P250 and P1K. Corresponding genes were linked to these DMCs chosen, and their LFCs were transformed by normal shrinkage. Data points are color-coded to indicate DEGs in red and non-DEGs in grey.

Further investigation focused on a set of 10,308 common DMCs, identified in at least one comparison. Distributions of methylation rates for these common DMCs exhibited a distinct left-skewed pattern across the exon, intron, P5K, and flanking regions (Fig. 5b). This observation strongly suggested that methylation differences resulting from altered spawning seasons primarily occurred within regions that were already highly methylated. In contrast, the P250 and P1K regions displayed nearly uniform methylation rates across a range of 0-100%, indicating significant methylation differences occurred irrespective of the underlying methylation rates.

These findings imply a potentially greater level of dynamic regulation of DNA methylation in P250 and P1K compared to the other regions.

### Filtering multiple DMCs identified key genes affected by spawning season alterations

We examined genes with multiple DMCs in three pairwise comparisons against the normal season. Since most genes had only a single DMC, especially in promoters (∼75% in P250 and ∼70% in P1K; Supplementary Figure S1), we avoided identifying differentially methylated regions (DMRs) using a sliding window approach. Instead, we filtered out genes strongly supported by multiple DMCs based on specific criteria (see Methods and Materials), resulting in a total of 11 genes with multiple DMCs on their promoters (Table 2; see the details in Supplementary Table S17).

**Table 2.**
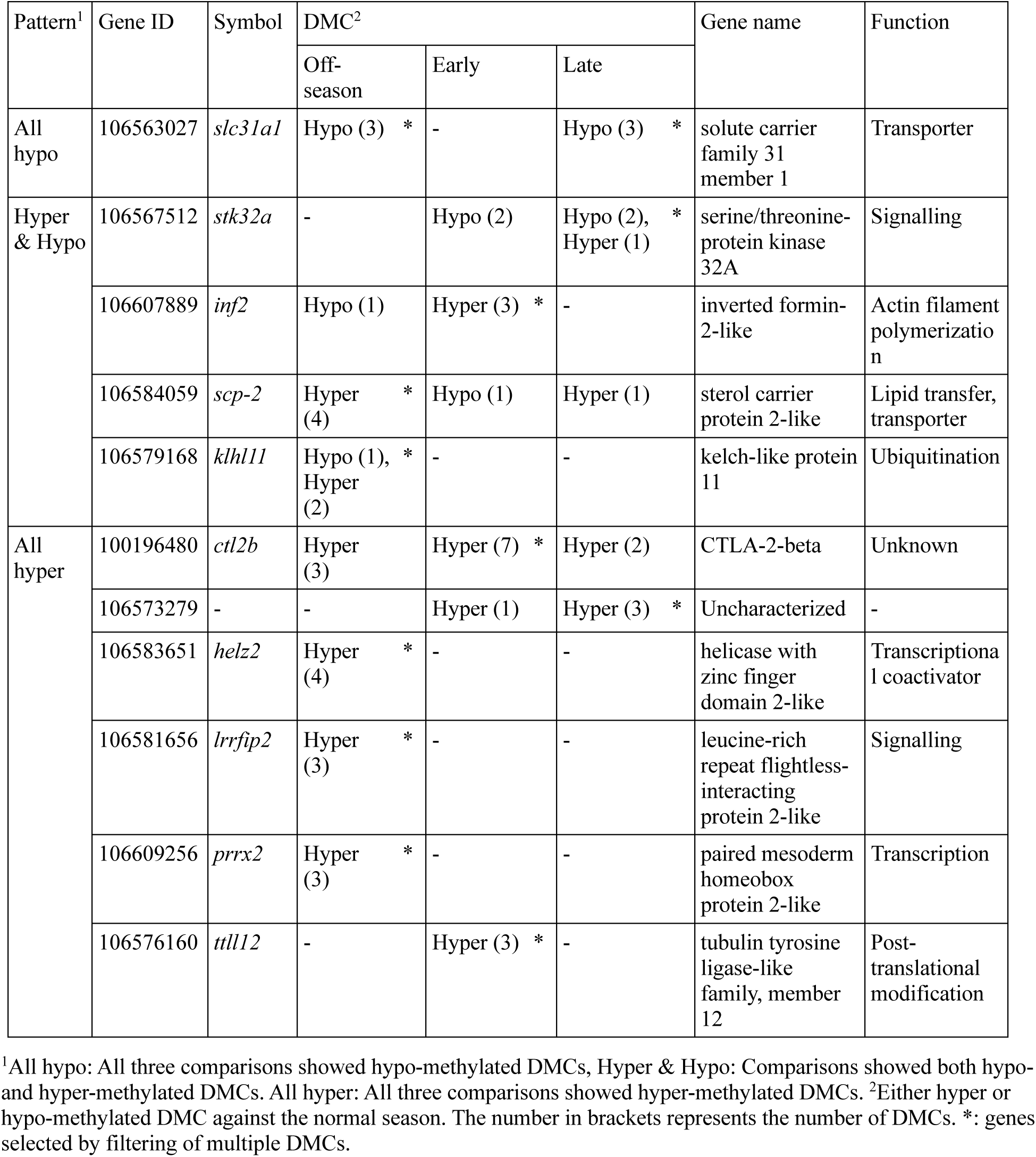
List of genes with multiple DMCs in their promoter region.

All 11 genes were identified by a single comparison except for *slc31a1*, which was identified in two comparisons (off-season vs. normal and late vs. normal). Most genes (9 out of 11) exhibited either hypo-methylated or hyper-methylated DMCs, but not both, within a single comparison (Table 2). Furthermore, approximately 60% of the genes (7 out of 11) consistently displayed either hypo-methylation or hyper-methylation even across multiple comparisons (’All hypo’ and ‘All hyper’ in the first column named ‘Patter’ of Table 2, respectively).

Literature analysis associated these genes with two biological functions: development (*inf2*) and information processing, which can be further classified into four sub categories as transporter (*slc31a1* and *scp-2*) (Stolowich et al., 2002; Zhou and Gitschier, 1997), signalling (*stk32a* and *lrrfip2*) (Alliance of Genome Resources, 2022; Bateman et al., 2022), post-translation (*klhl11* and *ttll12*) (Bateman et al., 2022; Brants et al., 2012), and transcription (*helz2* and *prrx2*) (Norris et al., 2000; Tomaru et al., 2006). However, *ctl2b* and an uncharacterised gene (Gene ID: 106573279) had no clear associations with specific biological functions. Although the DMCs on these genes were potentially linked to the underlying gene expression, none of the 11 genes were identified as DEGs in our differential gene expression analysis.

### Altered spawning seasons influenced gene expression and DNA methylation of cell cycle-related genes

To investigate potential associations between DNA methylation and gene expression, we combined DEGs and DMCs from three pairwise comparisons against the normal season (off-season vs. normal, early vs. normal, and late vs. normal) in P250 and P1K. This resulted in a total of seven DEGs that contain a total of 11 DMCs in their promoters (Fig. 5c, Table 3, Supplementary Table S18).

**Table 3.** List of DEGs that contain DMCs in their promoter region.

| Gene ID | Symbol | DMC<sup>1</sup> |  |  | DEG<sup>2</sup> |  |  | Gene name | Function |
| --- | --- | --- | --- | --- | --- | --- | --- | --- | --- |
|  |  | Off-season | Early | Late | Off-season | Early | Late |  |  |
| 106562317 | *caprin-1* | Hypo | (Hypo) | (Hypo) | Down | (Down) | (Down) | caprin-1 | Cell cycle |
| 106599887 | *cyp8b1* | Hypo | - | - | Down | - | - | 5-beta-cholestane-3-alpha,7-alpha-diol 12-alpha-hydroxylase | Metabolism, cytochrome P450 |
| 106588407 | *kifc1* | - | Hypo | Hypo | - | Down | Down | carboxy-terminal kinesin 2 | Cell cycle, Meiosis |
| 106582038 | *adrenodoxin* | - | Hypo |  | - | Down | - | adrenodoxin | Metabolism, cytochrome P450 |
| 106604665 | *slc43a1a* | - | Hypo | - | - | Up | (Up) | solute carrier family 43 member 1a | Transporter |
| 106561604 | *aurkb* | (Hyper) | Hyper | Hyper | - | Down | Down | aurora kinase B | Cell cycle |
| 106570052 | *lpin1* | - | Hyper | - | - | Up | - | phosphatidate phosphatase LPIN1 | Metabolism |

<sup>1</sup>Either hypo- or hyper-methylated DMC against the normal season. Parentheses indicate that the site was identified as DMC, but the corresponding genes were not identified as stringent DEG. <sup>2</sup>Either up- or down-regulation against the normal season. Parentheses indicate that down- or up-regulation was determined by non-stringent DEGs.

In addition to the significant results, we expanded the list of seven DEGs with additional information from all pair-wise comparisons (additional entries were marked with parentheses in Table 3). Among the identified genes, three (*caprin-1*, *kifc1*, and *aurkb*) exhibited consistent regulations of DMCs and DEGs compared to the normal season. Specifically, *caprin-1* and *kifc1* had hypo-methylated CpG sites, while *aurkb* had hyper-methylated CpG sites; all three genes showed down-regulated gene expression (Table 3). Notably, these three genes (*caprin-1*, *kifc1*, and *aurkb*) were associated with cell cycle regulation (Grill et al., 2004; Walczak et al., 1997; Wheatley et al., 2001), while the remaining genes were linked to metabolism (*cyp8b1*, *adrenodoxin*, and *lpin1*) (Grinberg et al., 2000; Hansson and Wikvall, 1982; Kanehisa and Goto, 2000a) and transport (*slc43a1a*) (Babu et al., 2003). These findings indicate that altered spawning seasons may have affected gene expression and epigenetic regulation of several genes associated with cell cycle control.

## Discussion

In the present study, we investigated the influence of altered spawning seasons on gene expression and DNA methylation in offspring. Previous studies emphasized the significant impacts of these changes on the nutritional status of both broodstock and their offspring, with no observed effects on the body weights of broodstock but in offspring (Skjærven et al., 2022; Skjaerven et al., 2020). Our primary aim was to reveal the transcriptomic and epigenetic effects resulting from observed variations by comparing liver samples from larvae at the first feeding stage.

### What insights do our omics analyses provide?

Gene expression analysis revealed distinct patterns associated with altered spawning seasons. Specifically, early and late seasons exhibited similar gene expression patterns when compared to the normal season. The differential expression of genes associated with metabolism, cellular processes, and organismal systems indicates that spawning seasons can have wide-ranging effects on biological pathways related to growth and development. Furthermore, the analysis with overlapped DEGs revealed that all three altered spawning seasons similarly affected genes associated with metabolism, tissue development, and information processing. Hence, while early and late seasons influenced a wide range of biological pathways in a similar manner, all three seasons, including the off-season, also similarly affected a certain set of genes, especially those strongly differentiated from the normal season.

DNA methylation analysis provided valuable insights into the epigenetic effects of different spawning seasons. Altered spawning seasons led to increased methylation rates for the early and normal seasons. Notably, these shifts primarily occurred in regions that were already highly methylated, but the rates were altered independently of the underlying methylation rates in promoter regions around transcription start sites. In addition, our findings across different developmental stages imply that methylation rates could serve as an indicator of age. However, more research is needed to understand the mechanism of methylation rate shifting and potential applications in aquaculture as well as age measure for stock assessment.

Several genes with multiple DMCs exhibited consistent hypo-or hyper-methylation patterns across multiple spawning seasons in promoter regions, suggesting a common epigenetic response to altered spawning seasons. These genes were associated with key biological functions, such as development and information processing, potentially indicating their role in growth performance. Although both gene expression and DNA methylation analyses resulted in similar biological functions impacted by altered spawning seasons, they showed little overlap between DEGs and genes with DMCs. This could be attributed to regulatory differences between gene expression and DNA methylation within the same biological pathway, or it might be a limitation caused by the restriction enzyme used in RRBS.

The observed correlations between DNA methylation and gene expression in our liver samples were weaker than anticipated. Note that the measurements were taken from siblings rather than the same individual due to the limited size of larvae livers. Notably, our findings also revealed inconsistent correspondence between hypo-methylation and active gene expression as well as between hyper-methylation and gene suppression. These results suggest that the regulation of DNA methylation is more intricate than commonly assumed as it is known that high promoter CpG methylation reduces gene expression, while low promoter CpG methylation allows for active gene expression (Lanata et al., 2018). Within promoter regions, not only DNA methylation but several other factors, including underlying gene expression levels, transcription factor binding sites, and chromatin remodelling, likely play a role in regulating actual gene expression.

### What are the significant biological differences observed in offspring?

#### Off-season vs. normal

Off-season spawning in June, facilitated through a RAS, resulted in Atlantic salmon offspring with lower larval weights compared to the normal and late seasons (Skjærven et al., 2022; Skjaerven et al., 2020). Offspring eggs also exhibited reduced levels of vitamin B12 and lipid classes relative to normal and late seasons (Skjærven et al., 2022; Skjaerven et al., 2020). Gene expression analysis indicated significant down-regulation of genes associated with central carbon metabolism (CCM). CCM involves several biological pathways, including the citric acid cycle, and acts as a major source of energy for growth and development. Moreover, lipids are interconnected with CCM through the utilisation of acetyl-CoA as a central metabolite (Westfall and Levin, 2018). Among the six genes with multiple DMCs from the off-season vs. normal comparison, sterol carrier protein2-like (*scp-2*) contained four hyper-methylated DMCs in its promoter. The *scp-2* gene encodes a transfer protein that plays a key role in intracellular lipid transport (Stolowich et al., 2002). Overall, RAS-based off-season spawning appeared to impact lipid-mediated regulations both transcriptionally and epigenetically. This negative impact on lipid-related mechanisms potentially accounted for impaired growth performance observed in body weight as our previous research indicated the importance of gene expression and epigenetic regulation of lipid regulation, specifically involving acetyl-CoA, for growth performance (Saito et al., 2021).

#### Early season vs. normal season

Offspring resulting from early season spawning in September also exhibited lower weights at the larval stage compared to those from normal and late seasons. However, the nutritional status of offspring eggs displayed higher levels of SAM/SAH, lysine, glutamine, and alanine (Skjærven et al., 2022; Skjaerven et al., 2020). Gene expression analysis revealed significant down-regulation of genes associated with cell cycle regulation, while genes involved in metabolism showed both down– and up-regulation. Among the five genes that exhibited differential expression and had at least one DMC in their promoters, two were associated with cell-cycle regulation. Specifically, the carboxy-terminal kinesin 2 (*kifc1*) gene promotes mitotic spindle assembly (Walczak et al., 1997), and the aurora kinase B (*aurkb*) gene is involved in mitotic progression (Wheatley et al., 2001). The *kifc1* gene was hypomethylated, while the *aurkb* gene was hypermethylated. Consequently, the early spawning season had an impact on cell-cycle regulation both transcriptionally and epigenetically. However, it is challenging to hypothesise that this negative impact on cell-cycle regulation was linked to impaired growth performance because similar effects were observed for the late season.

#### Late season vs. normal season

Late season spawning in January showed that offspring weights were almost the same, but slightly heavier than those from the normal season. The nutritional status of offspring eggs also exhibited similar levels of various nutrients and metabolites to the normal season, except for higher vitamin B12 and lower B-alanine (Skjærven et al., 2022). Gene expression patterns in the late season were comparable to those observed in the early season, showing significant down-regulation of genes related to cellular processes, especially cell cycle regulation. Two genes, *kifc1* and *aurkb*, showed differential expression and had at least one DMC in their promoters, and both were associated with cell-cycle regulation. Hence, the late season showed similar growth performance and nutritional status to the normal season, while the gene expression pattern was similar to the early season.

#### Three altered spawning seasons vs. normal season

Although strong similarities were observed in gene expression patterns between early and late spawning seasons, certain genes exhibited common regulation in both gene expression and DNA methylation across all three altered spawning seasons. Among the four DEGs identified in the off-season, choline-phosphate cytidylyltransferase B (*pcyt1b*) and bone morphogenetic protein 5-like (*bmp5*) showed concordant down-regulation across all three altered spawning seasons. The *pcyt1b* gene encodes an enzyme involved in phosphatidylcholine biosynthesis, playing essential roles in multiple metabolic pathways (Bateman et al., 2022), while the *bmp5* gene is involved in signalling (Kanehisa and Goto, 2000a). Additionally, all DMCs found in the promoter of the cytotoxic T lymphocyte-associated protein 2 beta (*ctl2b*) gene were hyper-methylated across all three altered spawning seasons. The function of *ctl2b* remains unknown, but it shares similarities with genes encoding cysteine proteinase, an enzyme involved in protein breakdown within cells (Denizot et al., 1989). Furthermore, among genes that were both differentially expressed and had at least one DMC in their promoters, the *caprin-1* gene exhibited consistent regulation, with all DMCs being hypo-methylated and its expression being suppressed. The *caprin-1* gene is associated with cell cycle regulation, playing a crucial role in cellular activation or proliferation (Grill et al., 2004). This indicates that the off-season also influenced cell cycle regulation to a certain degree.

### Why are our findings important in the context of aquaculture and nutritional studies?

Our study unveiled the significant effects of altered spawning season on offspring gene expression and DNA methylation patterns in Atlantic salmon. These findings provide valuable insights into assessing current and future growth potential, which can be challenging to determine using only nutritional status and growth performance measures. Understanding the gene regulation and epigenetic responses to altered spawning seasons will help develop strategies to optimize growth and production in aquaculture practices. Additionally, our study can be linked to the potential impact of increasing global ocean temperatures on the spawning of aquatic vertebrates, as well as the northward shift in habitats followed by changes in photoperiod, as both temperature and light are key abiotic factors influencing spawning seasons. Even though genetic and epigenetic profiles can be species– and even tissue-specific, the approaches we demonstrated here could be applied to other fish species whose spawning seasons are artificially altered in aquaculture practices. Nonetheless, further studies are needed to elucidate the fundamental effects caused by spawning season alterations across various species.

## Conclusions

The present study emphasises the substantial impact of altering spawning seasons on gene expression and DNA methylation patterns in Atlantic salmon offspring. Our observations indicate that RAS-based off-season spawning exerts an influence on lipid-mediated regulations in offspring, while sea-pen based early and late seasons affect the regulation of the offspring’s cellular processes, especially cell cycle regulation. These effects potentially play a crucial role in shaping both current and future growth performance.

These findings have important implications for aquaculture and nutritional studies, as they provide valuable tools for assessing growth potential and optimizing production strategies. Future research focusing on tissue and developmental stage-specific DNA methylation maps will enhance the accuracy of growth performance estimation and provide further insights into the gene regulation and epigenetic responses to altered spawning conditions.

## Acknowledgements

We thank Amelie Nemc and Bekir Ergüner for advice on RRBS analysis, and Hui-Shan Tung and Eva Mykkeltvedt for DNA and RNA extraction at the Institute of Marine Research (Bergen, Norway). This work was supported by the Norwegian Research Council under project no: 267787 (NutrEpi) and by the Institute of Marine Research under the Nutritional programming project.

## Author contributions

K.H.S conceived and designed the research. M.E. and M.M. conducted and designed the feeding trial. J.M.O.F. and C.B. prepared RNA-seq and RRBS data. T.S., M.E., and K.H.S. analysed and interpreted data. T.S. drafted the manuscript, and K.H.S revised it. The authors read and approved the final manuscript.

