## Supplementary information for "Altered spawning seasons of Atlantic salmon broodstock transcriptionally and epigenetically influence cell cycle and lipid-mediated regulations in their offspring"

### Tables

**Table S1**. Feeding regime overview for four spawning groups.

| | **Off-season** | **Early** | **Normal** | **Late** |
|---|---|---|---|---|
| **Starvation start date** | 03.03.2017 | 16.04.2017 | 26.07.2017 | 26.07.2017 |
| **Stripping date** | 20.06.2017 | 26.09.2017 | 23.11.2017 | 08.01.2018 |
| **Days of starvation** | 109 | 163 | 120 | 166 |
| **Egg/liter** | 6155 | 5835 | 5082 | 4613 |
| **Day° newly fertilized egg** | 2 | 2 | 2 | 2 |
| **Day° eye staged egg** | 364 | 363 | 362 | 359 |
| **Day° start feeding larvae** | 994 | 991 | 979 | 981 |

**Table S2**. Weights of broodstock and off-spring larvae in four spawning seasons.

| **Stage** | **Unit** | **Season** | **Weight mean** | **Weight SD** | **Tukey's HSD[1]** |
|---|---|---|---|---|---|
| Broodstock | kg | Off-season | 12.10 | 1.55 | - |
| | | Early | 11.52 | 1.02 | - |
| | | Normal | 11.74 | 0.84 | - |
| | | Late | 12.56 | 0.85 | - |
| Offspring | g | Off-season | 154.12 | 14.55 | a |
| | | Early | 142.04 | 15.52 | b |
| | | Normal | 184.37 | 20.42 | c |
| | | Late | 191.68 | 20.76 | d |

[1]Letters represent Tukey's HSD compact letter display after ANOVA tests ($p<0.05$).

**Table S3**. Nutritional status of the 1C metabolism in eggs.

| Nutrient | Unit | Season | Mean | SD | Tukey's HSD[1] |
|---|---|---|---|---|---|
| **Vit B6** | mg/kg ww | Off-season | 1.74 | 0.25 | - |
| | | Early | 1.56 | 0.05 | - |
| | | Normal | 1.76 | 0.18 | - |
| | | Late | 1.81 | 0.19 | - |
| **Folate** | mg/kg ww | Off-season | 1.64 | 0.12 | ab |
| | | Early | 1.26 | 0.14 | a |
| | | Normal | 1.82 | 0.25 | b |
| | | Late | 1.60 | 0.25 | ab |
| **Vit B12** | µg/100 g ww | Off-season | 62.84 | 10.94 | a |
| | | Early | 108.34 | 7.32 | b |
| | | Normal | 107.49 | 19.20 | b |
| | | Late | 166.76 | 8.25 | c |
| **SAM/SAH** | - | Off-season | 3.70 | 0.49 | a |
| | | Early | 4.72 | 0.29 | b |
| | | Normal | 3.17 | 0.36 | ac |
| | | Late | 2.71 | 0.34 | c |

[1]Letters represent Tukey's HSD compact letter display after ANOVA tests ($p<0.05$).

**Table S4**. Nutritional status of lipid classes in eggs.

| Nutrient | Unit | Season | Mean | SD | Tukey's HSD[1] |
|---|---|---|---|---|---|
| **Cholesterol** | mg/g ww | Off-season | 3.20 | 0.67 | a |
| | | Early | 5.76 | 0.59 | b |
| | | Normal | 5.80 | 0.22 | b |
| | | Late | 5.78 | 0.08 | b |
| **Sum of lipids** | mg/g ww | Off-season | 97.52 | 17.33 | a |
| | | Early | 167.80 | 8.87 | b |
| | | Normal | 173.40 | 14.81 | b |
| | | Late | 167.60 | 9.66 | b |

[1]Letters represent Tukey's HSD compact letter display after ANOVA tests ($p<0.05$).

**Table S5**. Nutritional status of the citric acid cycle in eggs.

| Nutrient | Unit | Season | Mean | SD | Tukey's HSD[1] |
|---|---|---|---|---|---|
| **Glutamate** | µmol/g ww | Off-season | 3.44 | 0.69 | a |
| | | Early | 4.08 | 0.30 | a |
| | | Normal | 1.87 | 0.43 | b |
| | | Late | 2.15 | 0.51 | b |
| **L-Lysine** | µmol/g ww | Off-season | 0.62 | 0.17 | a |
| | | Early | 0.89 | 0.10 | b |
| | | Normal | 0.53 | 0.10 | a |
| | | Late | 0.49 | 0.03 | a |

[1]Letters represent Tukey's HSD compact letter display after ANOVA tests ($p<0.05$).

**Table S6**. Nutritional status of the Cahill cycle in eggs.

| Nutrient | Unit | Season | Mean | SD | Tukey's HSD[1] |
|---|---|---|---|---|---|
| **L-Alanine** | µmol/g ww | Off-season | 1.53 | 0.47 | a |
| | | Early | 2.05 | 0.20 | b |
| | | Normal | 1.39 | 0.14 | a |
| | | Late | 1.57 | 0.06 | ab |
| **L-Glutamine** | µmol/g ww | Off-season | 1.16 | 0.26 | a |
| | | Early | 1.61 | 0.11 | b |
| | | Normal | 0.79 | 0.06 | c |
| | | Late | 0.87 | 0.12 | c |
| **Urea** | µmol/g ww | Off-season | 0.21 | 0.08 | a |
| | | Early | 0.28 | 0.08 | ab |
| | | Normal | 0.38 | 0.11 | bc |
| | | Late | 0.50 | 0.09 | c |
| **B-Alanine** | µmol/g ww | Off-season | 0.05 | 0.01 | ab |
| | | Early | 0.10 | 0.03 | c |
| | | Normal | 0.07 | 0.01 | b |
| | | Late | 0.03 | 0.01 | a |

[1]Letters represent Tukey's HSD compact letter display after ANOVA tests ($p<0.05$).

**Table S7**. List of RNA-seq samples with alignment and mapping statistics.

| Season | Sample | M Seqs[1] | M Aligned[2] | % Aligned | M Assigned[3] | % Assigned |
|---|---|---|---|---|---|---|
| **Off-season** | O1 | 44.5 | 37.4 | 84.0% | 33.1 | 61.3% |
| | O2 | 46.4 | 38.8 | 83.7% | 33.8 | 60.0% |
| | O3 | 46.2 | 38.6 | 83.5% | 33.6 | 60.0% |
| | O4 | 46.6 | 39.2 | 84.1% | 34.4 | 61.3% |
| | O5 | 35.6 | 29.8 | 83.8% | 26.7 | 61.6% |
| **Early** | E1 | 44.5 | 37.7 | 84.8% | 32.9 | 61.6% |
| | E2 | 42.9 | 36.4 | 84.8% | 32.1 | 62.6% |
| | E3 | 41.1 | 34.7 | 84.3% | 30.8 | 62.7% |
| | E4 | 36.3 | 30.5 | 83.8% | 27.3 | 62.2% |
| | E5 | 36.2 | 30.6 | 84.5% | 27 | 61.4% |
| **Normal** | N1 | 33.2 | 27.8 | 83.7% | 25 | 62.0% |
| | N2 | 43.7 | 36.4 | 83.2% | 32.8 | 63.4% |
| | N3 | 44 | 36.9 | 84.0% | 32.9 | 62.0% |
| | N4 | 39.8 | 33.6 | 84.6% | 29.7 | 62.0% |
| | N5 | 37.1 | 31.3 | 84.4% | 28.2 | 62.9% |
| **Late** | L1 | 37.2 | 31.5 | 84.9% | 27.5 | 62.0% |
| | L2 | 47.4 | 39.9 | 84.2% | 35.7 | 62.4% |
| | L3 | 41.9 | 35.3 | 84.2% | 31.3 | 62.1% |
| | L4 | 43.3 | 36.5 | 84.3% | 32.5 | 62.1% |
| | L5 | 44.4 | 37.5 | 84.5% | 33.1 | 61.9% |

[1]Number of reads in millions. [2]Number of uniquely aligned reads to the reference genome. [3]Number of uniquely mapped reads to genes for quantification.

**Table S8**. One enriched KEGG pathway identified by ORA for off-season vs. normal DEGs.

| ID | Enriched KEGG pathway | GeneRatio | BgRatio | P-value | Adjusted p-value |
|---|---|---|---|---|---|
| **sasa01200** | Carbon metabolism | 5/44 | 247/11637 | 2.29E-03 | 4.58E-02 |

**Table S9**. Six enriched KEGG pathways identified by ORA for early vs. normal DEGs.

| ID | Enriched KEGG pathway | GeneRatio | BgRatio | P-value | Adjusted p-value |
|---|---|---|---|---|---|
| **sasa04110** | Cell cycle | 16/104 | 275/11546 | 2.81E-09 | 2.39E-07 |
| **sasa04914** | Progesterone-mediated oocyte maturation | 9/104 | 204/11546 | 9.07E-05 | 3.86E-03 |
| **sasa04115** | p53 signaling pathway | 7/104 | 147/11546 | 3.56E-04 | 9.63E-03 |
| **sasa04114** | Oocyte meiosis | 9/104 | 253/11546 | 4.53E-04 | 9.63E-03 |
| **sasa03320** | PPAR signaling pathway | 6/104 | 137/11546 | 1.47E-03 | 2.49E-02 |
| **sasa00330** | Arginine and proline metabolism | 5/104 | 99/11546 | 1.97E-03 | 2.79E-02 |

**Table S10**. Nine enriched KEGG pathways by ORA for late vs. normal DEGs.

| ID | Enriched KEGG pathway | GeneRatio | BgRatio | P-value | Adjusted p-value |
|---|---|---|---|---|---|
| **sasa04110** | Cell cycle | 13/91 | 278/11587 | 2.38E-07 | 2.14E-05 |
| **sasa04914** | Progesterone-mediated oocyte maturation | 9/91 | 204/11587 | 3.07E-05 | 1.38E-03 |
| **sasa04115** | p53 signaling pathway | 7/91 | 148/11587 | 1.59E-04 | 3.63E-03 |
| **sasa04114** | Oocyte meiosis | 9/91 | 253/11587 | 1.61E-04 | 3.63E-03 |
| **sasa00230** | Purine metabolism | 9/91 | 269/11587 | 2.55E-04 | 4.59E-03 |
| **sasa00480** | Glutathione metabolism | 5/91 | 90/11587 | 6.95E-04 | 1.04E-02 |
| **sasa00983** | Drug metabolism - other enzymes | 5/91 | 102/11587 | 1.22E-03 | 1.57E-02 |
| **sasa04218** | Cellular senescence | 9/91 | 375/11587 | 2.67E-03 | 2.41E-02 |
| **sasa03320** | PPAR signaling pathway | 5/91 | 137/11587 | 4.43E-03 | 3.63E-02 |

**Table S11**. Top 5 DEGs identified at least two pairwise comparisons.

| Comp[1] | Gene ID | LFC | Adjusted[2] | Symbol[3] | Gene name |
|---|---|---|---|---|---|
| **Off-season** | 106612264 | 5.22 | 6.04E-04 | - | - |
| | 106563634 | -1.87 | 3.54E-03 | pcyt1b | choline-phosphate cytidylyltransferase B |
| | 106601246 | 1.85 | 5.52E-03 | (cbx7) | (Chromobox 7) |
| | 106612672 | -1.20 | 1.34E-02 | bmp5 | bone morphogenetic protein 5-like |
| | 106592454 | -1.70 | 3.59E-02 | pc | pyruvate carboxylase, mitochondrial-like |
| **Early** | 106608456 | -2.28 | 9.69E-16 | prox3 | prospero homeobox 3 |
| | 106613828 | 3.01 | 9.27E-15 | crim1 | cysteine-rich motor neuron 1 protein-like |
| | 106566973 | 1.27 | 2.83E-09 | krt8 | keratin, type II cytoskeletal 8 |
| | 106585803 | -1.12 | 1.39E-08 | rhobtb4 | Rho related BTB domain containing 4 |
| | 106584736 | -3.34 | 1.14E-07 | tcf12 | transcription factor 12 |
| **Late** | 106583637 | 3.85 | 6.75E-17 | klf15 | Krueppel-like factor 15 |
| | 106608456 | -1.74 | 3.44E-09 | prox3 | prospero homeobox 3 |
| | 106568312 | -2.63 | 4.18E-09 | pitpnm2 | membrane-associated phosphatidylinositol transfer protein 2-like |
| | 106613828 | 3.02 | 1.32E-08 | crim1 | cysteine-rich motor neuron 1 protein-like |
| | 106569008 | 2.93 | 1.36E-07 | cyp7a1 | cytochrome P450 7A1 |

[1]Pair-wise comparisons against the normal season. [2]Adjusted p-value. [3]When gene symbols were unavailable, we utilised the gene symbols from either the latest NCBI version or UniProt as substitutes. The gene symbol of 106601246 (cbx7) was estimated by OrthoDB.

**Table S12**. List of RRBS samples with alignment and methylation calling statistics.

| Season | Sample | M Seqs[1] | % Aligned[2] | % mCpG[3] | % mCHG[4] | % mCHH[5] | M C's[6] |
|---|---|---|---|---|---|---|---|
| **Off-season** | O1 | 56.4 | 49.3% | 80.6% | 1.0% | 0.9% | 275.8 |
| | O2 | 72.1 | 49.4% | 80.9% | 1.0% | 0.9% | 353.1 |
| | O3 | 45.6 | 49.1% | 80.9% | 1.0% | 0.9% | 220.3 |
| | O4 | 57.1 | 49.3% | 80.5% | 1.0% | 0.9% | 276 |
| | O5 | 49.2 | 48.6% | 80.9% | 1.1% | 1.0% | 230.2 |
| **Early** | E1 | 65.2 | 46.4% | 82.0% | 1.0% | 0.9% | 300 |
| | E2 | 73.3 | 48.9% | 81.1% | 1.0% | 0.9% | 348.1 |
| | E3 | 59.8 | 49.5% | 80.8% | 1.0% | 0.9% | 294.7 |
| | E4 | 45.6 | 49.2% | 80.4% | 1.0% | 0.9% | 217.6 |
| | E5 | 53 | 50.0% | 80.0% | 1.0% | 0.9% | 262.2 |
| **Normal** | R1 | 51 | 50.9% | 79.7% | 1.0% | 0.9% | 254.5 |
| | R2 | 50.1 | 50.4% | 80.8% | 1.0% | 0.9% | 246.6 |
| | R3 | 60.6 | 49.0% | 80.4% | 1.0% | 0.9% | 291.6 |
| | R4 | 49.4 | 49.2% | 80.9% | 1.0% | 0.9% | 240.4 |
| | R5 | 55 | 49.5% | 81.6% | 1.0% | 0.9% | 265.5 |
| **Late** | L1 | 42.3 | 49.0% | 80.6% | 1.0% | 0.9% | 202.7 |
| | L2 | 57.9 | 48.6% | 81.1% | 1.1% | 1.0% | 274.1 |
| | L3 | 60 | 48.5% | 80.8% | 1.0% | 0.9% | 286.1 |
| | L4 | 44 | 48.9% | 80.4% | 1.0% | 0.9% | 210.6 |
| | L5 | 43.5 | 49.0% | 81.2% | 1.0% | 0.9% | 213 |

[1]Number of reads in millions. [2]Number of uniquely aligned reads to the reference genome. [3,4,5]Percentage of methylated cytosine in CpG, CHG, and CHH sites, respectively where H represents non-cytosine. [6]Count of total CpG sites with methylation calling information in millions.

**Table S13**. Comparisons of methylation raters between treatment groups in three life stages.

| Stage | Group1 | Group2 | Adjusted p-value[1] | Significant[2] |
|---|---|---|---|---|
| **Larvae** | Early | Off-season | 0.37 | |
| | Normal | Off-season | 0.37 | |
| | Late | Off-season | 0.37 | |
| | Normal | Early | 0.37 | |
| | Late | Early | 0.55 | |
| | Late | Normal | 0.37 | |
| **Post-smolt** | 1C+ | Ctrl | 2.47E-04 | *** |
| | 1C++ | Ctrl | 1.85E-03 | * |
| | 1C++ | 1C+ | 0.19 | |
| **Harvest** | L2 | L1 | 0.95 | |
| | L3 | L1 | 0.95 | |
| | L3 | L2 | 0.95 | |

[1]P-values were adjusted by Holm. [2]Distributions were significantly different ('*': adjusted p-value<0.05, '**': adjusted p-value<0.01, "***': adjusted p-value<0.001).

**Table S14**. Comparisons of methylation raters between life stages by genomic regions.

| Region | Group1 | Group2 | Adjusted p-value[1] | Significant[2] |
|---|---|---|---:|---|
| **Exon** | Post-smolt | Larvae | 1.07E-04 | *** |
| | Harvest | Larvae | 2.23E-11 | *** |
| | Harvest | Post-smolt | 4.15E-07 | *** |
| **Intron** | Post-smolt | Larvae | 1.97E-02 | * |
| | Harvest | Larvae | 2.23E-11 | *** |
| | Harvest | Post-smolt | 3.82E-09 | *** |
| **P250** | Post-smolt | Larvae | 0.92 | |
| | Harvest | Larvae | 0.92 | |
| | Harvest | Post-smolt | 0.92 | |
| **P1K** | Post-smolt | Larvae | 6.15E-13 | *** |
| | Harvest | Larvae | 1.11E-11 | *** |
| | Harvest | Post-smolt | 0.26 | |
| **P5K** | Post-smolt | Larvae | 6.15E-13 | *** |
| | Harvest | Larvae | 1.11E-11 | *** |
| | Harvest | Post-smolt | 9.05E-08 | *** |
| **Flank** | Post-smolt | Larvae | 1.38E-11 | *** |
| | Harvest | Larvae | 1.38E-11 | *** |
| | Harvest | Post-smolt | 2.69E-06 | *** |
| **Misc** | Post-smolt | Larvae | 6.15E-13 | *** |
| | Harvest | Larvae | 2.23E-10 | *** |
| | Harvest | Post-smolt | 4.03E-12 | *** |
| **IGR** | Post-smolt | Larvae | 1.56E-03 | ** |
| | Harvest | Larvae | 1.56E-03 | ** |
| | Harvest | Post-smolt | 8.08E-08 | *** |

[1]P-values were adjusted by Holm. [2]Distributions were significantly different ('*': adjusted p-value<0.05, '**': adjusted p-value<0.01, "***': adjusted p-value<0.001).

**Table S15**. Comparisons of methylation raters between spawning seasons by genomic regions.

| Region | Group1 | Group2 | Adjusted p-value[1] | Significant[2] |
|---|---|---|---|---|
| **Exon** | Early | Off-season | 0.62 | |
| | Normal | Off-season | 0.84 | |
| | Late | Off-season | 0.66 | |
| | Normal | Early | 0.62 | |
| | Late | Early | 0.62 | |
| | Late | Normal | 0.66 | |
| **Intron** | Early | Off-season | 0.30 | |
| | Normal | Off-season | 0.30 | |
| | Late | Off-season | 0.84 | |
| | Normal | Early | 0.66 | |
| | Late | Early | 0.33 | |
| | Late | Normal | 0.30 | |
| **P250** | Early | Off-season | 1 | |
| | Normal | Off-season | 1 | |
| | Late | Off-season | 1 | |
| | Normal | Early | 1 | |
| | Late | Early | 1 | |
| | Late | Normal | 1 | |
| **P1K** | Early | Off-season | 1 | |
| | Normal | Off-season | 1 | |
| | Late | Off-season | 1 | |
| | Normal | Early | 1 | |
| | Late | Early | 1 | |
| | Late | Normal | 1 | |
| **P5K** | Early | Off-season | 0.66 | |
| | Normal | Off-season | 0.62 | |
| | Late | Off-season | 0.84 | |
| | Normal | Early | 0.63 | |
| | Late | Early | 0.62 | |
| | Late | Normal | 0.62 | |
| **Flank** | Early | Off-season | 0.29 | |
| | Normal | Off-season | 0.63 | |
| | Late | Off-season | 1 | |
| | Normal | Early | 1 | |
| | Late | Early | 0.29 | |
| | Late | Normal | 0.62 | |
| **Misc** | Early | Off-season | 0.84 | |
| | Normal | Off-season | 0.84 | |
| | Late | Off-season | 0.84 | |
| | Normal | Early | 0.84 | |
| | Late | Early | 0.84 | |
| | Late | Normal | 0.84 | |
| **IGR** | Early | Off-season | 0.83 | |
| | Normal | Off-season | 0.83 | |
| | Late | Off-season | 0.83 | |
| | Normal | Early | 0.83 | |
| | Late | Early | 0.83 | |
| | Late | Normal | 1 | |

[1]P-values were adjusted by Holm. [2]Distributions were significantly different ('*': adjusted p-value<0.05, '**': adjusted p-value<0.01, "***': adjusted p-value<0.001).

**Table S16**. Average methylation raters of spawning seasons by genomic regions.

| Region | Methylation rate (%) | | | | Order |
|---|---|---|---|---|---|
| | Off-season | Early | Normal | Late | |
| **Exon** | 74.590 | 74.965 | 74.654 | 74.784 | Off-season < Normal < Late < Early |
| **Intron** | 85.142 | 85.412 | 85.524 | 85.193 | Off-season < Late < Early < Normal |
| **P250** | 21.923 | 22.299 | 22.087 | 21.755 | Late < Off-season < Normal < Early |
| **P1K** | 44.358 | 44.507 | 44.910 | 44.595 | Off-season < Early < Late < Normal |
| **P5K** | 76.011 | 76.230 | 76.363 | 75.994 | Late < Off-season < Early < Normal |
| **Flank** | 78.510 | 78.777 | 78.674 | 78.418 | Late < Off-season < Normal < Early |
| **Misc** | 83.891 | 84.232 | 84.320 | 84.099 | Off-season < Late < Early < Normal |
| **IGR** | 87.327 | 87.523 | 87.634 | 87.595 | Off-season < Late < Early < Normal |

**Table S17**. List of genes with multiple DMCs in P250 and P1K between spawning seasons.

| **Comp**[1] | **Gene ID** | **#DMCs (P250)**[2] | | | **#DMCs (P1K)**[3] | | | **Gene symbol**[4] | **Gene name** |
|---|---|---|---|---|---|---|---|---|---|
| | | Total | Hypo | Hyper | Total | Hypo | Hyper | | |
| **Off-season** | 106579168 | **3** | 1 | 2 | 0 | 0 | 0 | *klhl11* | kelch-like protein 11 |
| | 106609256 | **3** | 0 | 3 | 0 | 0 | 0 | *prrx2* | paired mesoderm homeobox protein 2-like |
| | 106583651 | 0 | 0 | 0 | **4** | 0 | 4 | *helz2* | helicase with zinc finger domain 2-like |
| | 106584059 | 0 | 0 | 0 | **4** | 0 | 4 | *scp-2* | sterol carrier protein 2-like |
| | 106563027 | 0 | 0 | 0 | **3** | 3 | 0 | *slc31a1* | solute carrier family 31 member 1 |
| | 106581656 | 0 | 0 | 0 | **3** | 0 | 3 | *lrrfip2* | leucine-rich repeat flightless-interacting protein 2-like |
| **Early** | 106607889 | **3** | 0 | 3 | 0 | 0 | 0 | *inf2* | inverted formin-2-like |
| | 100196480 | 3 | 0 | 3 | **4** | 0 | 4 | *ctl2b* | CTLA-2-beta |
| | 106576160 | 0 | 0 | 0 | **3** | 0 | 3 | *ttll12* | tubulin tyrosine ligase-like family, member 12 |
| **Late** | 106567512 | 0 | 0 | 0 | **3** | 2 | 1 | *stk32a* | serine/threonine-protein kinase 32A |
| | 106563027 | 0 | 0 | 0 | **3** | 3 | 0 | *slc31a1* | solute carrier family 31 member 1 |
| | 106573279 | 0 | 0 | 0 | **3** | 0 | 3 | - | Uncharacterized |

[1]Pair-wise comparisons against the normal season. [2,3]Number of DMCs in P250 and P1K regions. Genes were selected when they had three or more DMCs either in P250 or P1K. Counts in bold face indicates genes with three or more DMCs. [4]When gene symbols were unavailable, we utilised the gene symbols from either the latest NCBI version or UniProt as substitutes.

**Table S18.** List of DEGs with at least one DMC in P250 and P1K between spawning seasons.

| Cmp[1] | Gene ID | LFC | Chr[2] | Start[3] | Reg | Mdf[4] | Symbol | Gene name[5] |
|---|---|---|---|---|---|---|---|---|
| **Off-season** | 106562317 | -0.72 | NC_027310 | 24493341 | P250 | -21.97 | *caprin-1* | caprin-1 |
| | 106599887 | -0.69 | NC_027302 | 32167196 | P250 | -18.86 | *cyp8b1* | 5-beta-cholestane-3-alpha,7-alpha-diol 12-alpha-hydroxylase |
| **Early** | 106561604 | -0.85 | NC_027309 | 105304013 | P1K | 23.04 | *aurkb* | aurora kinase B |
| | 106570052 | 1.04 | NC_027300 | 15370776 | P1K | 17.95 | *lpin1* | phosphatidate phosphatase LPIN1 |
| | 106582038 | -0.85 | NC_027320 | 29797092 | P1K | -16.58 | *adrenodoxin* | adrenodoxin |
| | 106600949 (*) | 1.09 | NC_027302 | 52643520 | P1K | -18.40 | - | ferritin, middle subunit |
| | | | NC_027302 | 52643563 | P1K | -26.45 | | |
| | 106604665 | 0.99 | NC_027304 | 30317427 | P1K | -18.38 | *slc43a1a* | solute carrier family 43 member 1a |
| | 106588407 | -1.00 | NC_027326 | 9996221 | P250 | -27.36 | *caprin-1* | caprin-1 |
| **Late** | 106561604 | -0.73 | NC_027309 | 105304013 | P1K | 25.66 | *aurkb* | aurora kinase B |
| | 106588407 | -0.93 | NC_027326 | 9996221 | P250 | -24.39 | *kifc1* | carboxy-terminal kinesin 2 |

*: Gene ID 106600949 is discontinued in the current NCBI as of 2023. Hence, it is excluded from the main text.

[1]Pair-wise comparisons against the normal season. [2,3]Chromsome (Chr) and start position (Start) of DMCs. Suffix of chromosome ".1" was removed. For instance, the full name of NC_027310 is NC027310.01. [4]Mdf: methylation rate difference of DMC. [5]When gene symbols were unavailable, we utilised the gene symbols from either the latest NCBI version or UniProt as substitutes.

### Figures

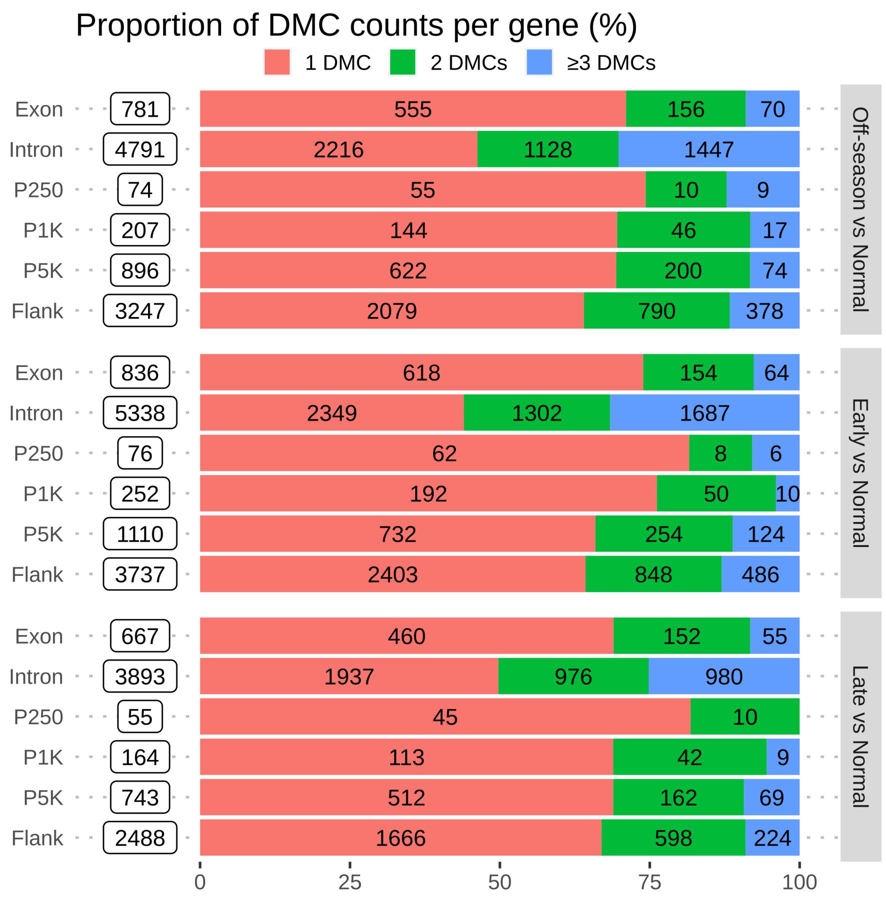


**Figure S1.** DMC counts per gene by three pairwise comparisons.

Stacked barplots showing the number of DMCs per gene, 1 DMC (red), 2 DMCs (green), and greater than or equal to 3 DMCs (blue), in six different genomic regions. The x-axis represents proportions of DMC counts in percentage.
